# Tip-links serve as force-pass filter to fulfil the role of gating-springs

**DOI:** 10.1101/2022.08.10.503460

**Authors:** Nisha Arora, Jagadish P. Hazra, Sandip Roy, Gaurav K. Bhati, Sarika Gupta, Abhishek Chaudhuri, Amin Sagar, Sabyasachi Rakshit

## Abstract

Tip-links as gating-spring in the mechanotransduction in hearing is still a debate. While the molecular elasticity of individual tip-link proteins warrants its candidature, the apparent rigidity from the heterotetrameric tip-links assembly refutes the claim. Using force-clamp experiments and simulations, we report that the heterotetrameric assembly is the natural selection for the gating-springs. Tip-links follow slip-ideal-slip bonds with increasing force. While in slip, the complex dissociates monotonously, ideal-bond interface responds indifferently to various auditory inputs. Insensitivity to forces renders tip-links as low-force pass filter, characteristic of gating-spring. Individual tip-links, however, forms slip-catch-slip bonds under tension. While catch bonds turn stronger with force from loud sound, our Langevin dynamics indicated the transition from slip-catch to slip-ideal bonds as cooperative effect of the dimers of individual protein complexes in tip-links. From molecular dynamics, we deciphered the molecular mechanism of catch bonds and its importance in deafness.

## Introduction

Hearing is one of the well-developed sensory processes in our body. Sound as stimuli induce oscillations in the inner ear fluid and deflect hair bundles, called stereocilia, protruding from the apical end of hair cells(Hudspeth, 1997). The deflection of stereocilia inflicts a shear tension on tip-links, long filamentous protein complexes, that connect two adjacent stereocilia in the direction of fluid-propagation(Pickles et al., 1984a). Tip-links convey the tension to mechanoelectrical transduction (MET) channel for response. The amplitude of the positive deflection of stereocilia depends on the intensity of input sound(Corey and Hudspeth, 1983), and so as the tension generated at the tip-link region. Thus, the physical presence of a gating-spring was realized in the mechano-transduction process to discriminate the tension from various auditory inputs, convey a threshold force to trigger MET channel opening, and damp overexcitation (Hudspeth, 1989).

Tip-links, a doublet of cadherin couples, was conceptualized as the simplest gating-spring in hearing(Basu et al., 2016; Kazmierczak et al., 2007; Pickles et al., 1984). However, the conformational topography from electron microscopy and all-atom simulations projected tip-links as too rigid to be gating-spring(Kachar et al., 2000; Sotomayor et al., 2010). It was proposed that additional molecular element like ankyrin with large viscoelasticity that are attached with tip-links in series may form an extended tip-links complex and serve as gating-spring (Sotomayor et al., 2005). Counter-arguably, more viscoelastic elements require longer time for relaxation and may defeat the limit of faster relaxation of 10 μs or less at high auditory frequencies(Corey and Hudspeth, 1983; Muller and Gillespie, 2010). Moreover, a simplest gating-spring functional for a wide range of auditory intensity decibels and frequencies yet with minimal molecular elements is naturally preferred in the MET in hearing. Thus, conceptually and casuistically, tip-links is still the most preferred gating-springs. However, it demands further clarity in the functional model to establish tip-links as gating-springs. Traditionally, temporal response to constant force or stress has been the classical physics experiment to study the viscoelasticity of soft matters, extrapolation to which provided the spring models. Notably, several pulling experiments have already been done with individual tip-links proteins which deciphered the axial elasticity of the proteins(Arora et al., 2021; Bartsch et al., 2018; Oroz et al., 2019). While these studies quantitatively identify and characterize the elastic and non-elastic components of tip-links proteins, they failed to identify the characteristic features of a gating-spring. Using single-molecule experiments and detailed simulations, we expose the tip-links under constant tensile forces and show how the temporal force-response of the binding interface reflects the dynamic modulation of external forces by tip-links as gating springs.

Tip-links is formed by the heteromeric interaction of two homodimeric non-classical cadherin family proteins, Cadherin-23 (Cdh23) and Protocadherin-15 (Pcdh15)(Kazmierczak et al., 2007). It connects to MET channel directly. Both proteins, like their other family members, possess a cytosolic region, a transmembrane part, and an extracellular (EC) part. Chemically, all three regions of the Cdh23 and Pcdh15 proteins differ from each other. The lengths of the EC regions are different too; Cdh23 has 27 EC domains and Pcdh15 has only 11. However, topologically all the domains are similar across tip-links. Tip-links complex is mediated by the interactions between two outermost domains (EC1-2) of Cdh23 and Pcdh15 in handshake-conformation(Sotomayor et al., 2012)(**Figure 1a,b**). The rest of the domains of tip-links cadherins facilitate lateral interactions and may form cis-homodimers(Honig et al., 2018; Jaiganesh et al., 2018). Apart from EC domains, there are linkers connecting two neighbouring domains. Linkers introduce kinks and bents in cadherin(Araya-Secchi et al., 2016; Powers et al., 2017). Kinks and bents in proteins are mechanical linkages that contribute to the elasticity of the tertiary structure(Granzier et al., 1997; Lee et al., 2006). Accordingly, individual tip-links proteins and the tip-links complex too can exploit their tertiary structure elasticity in response to mechanical stimuli and contribute to force-dissipation(Arora et al., 2021; Bartsch et al., 2018; Choudhary et al., 2020; Oroz et al., 2019). In fact, the mechanical origin of force-dissipation for individual tip-links is previously identified from non-equilibrium simulations and experiments (Arora et al., Choudhary et al., 2020), however, their impact on the tip-links complex interface is still elusive.

**Figure 1:**
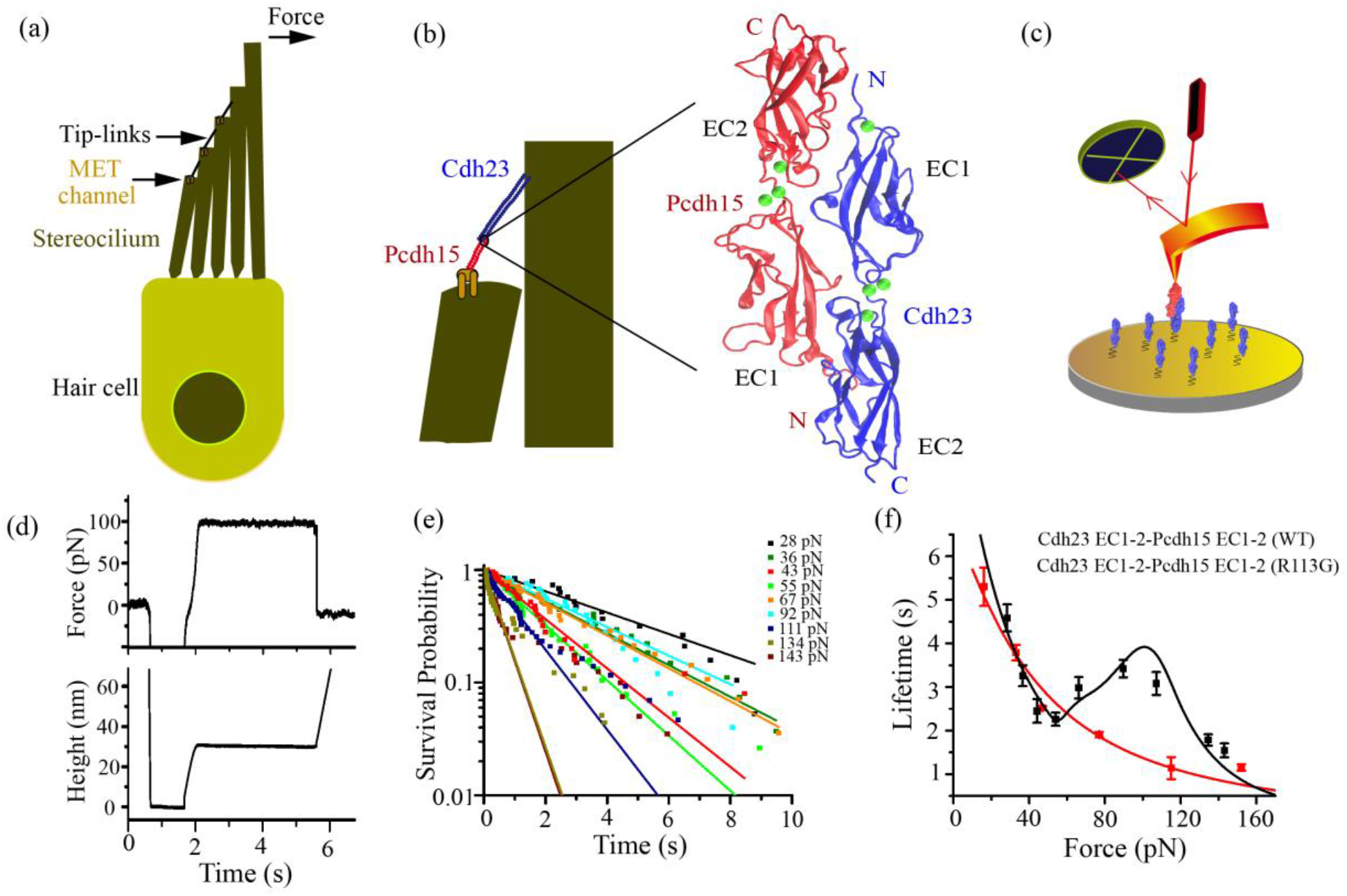
Tip-links follow slip-catch-slip bonds under tension. **(a)** Schematic of a hair-cell with hair-bundle at the apical end where **e**ach stereocilium in the hair bundle is connected to the next taller stereocilium through a filamentous element at the tips, known as tip-links. **(b)** The interacting domains in tip-links are highlighted. A zoomed region depicts the ribbon diagram of a tip-link complex interface, mediated by the two outermost domains of the constituent proteins, Cdh23 from top and Pcdh15 from base. **(c)** Schematic of the AFM setup used for the force-clamp experiment. Pcdh15 EC 1-2 (red) is shown at the cantilever and Cdh23 EC1-2 (blue) is onto the surface. **(d)** Typical force vs. time (top) and height vs. time (bottom) graphs, respectively obtained in an AFM force-clamp experiment. In this example, clamping is done at 100 pN. The bond-survival time is measured from the duration of clamp-time. **(e)** the bond-survival probabilities for the tip-links complex are shown here for the nine clamping forces. The solid lines represent the exponential-decay fit. We obtained the bond-lifetime at every force from the fitted curve. **(f)** The dependence of bond-lifetime with clamping forces are shown here for wild-type (black squares) and R113G-mutant complex (red squares). The wild-type (WT) tip-links complex shows a tri-phasic slip-catch-slip behaviour. Error bars are the standard errors obtained from the exponential fitting of the survival plots. Black solid line is the force-induced stronger-binding kinetic model fit. Red solid line is the Bell model fit.

For homo sapiens, the perceived range of sound is 20 Hz-20 kHz in frequency and 5-120 dB in intensity, referring to a tension of 10-100 pN on tip-links loading at kHz rate(Hudspeth, 1992). However, the average lifetime of individual tip-links under resting force (∼10 pN) is ∼8 s(Mulhall et al., 2021). Moreover, the lifetime of tip-links drops exponentially with force when measured at slower force-loading rates(Mulhall et al., 2021), indicating a nearly disappearing existence of tip-links in a nightclub or orchestra. It is thus intriguing to decipher the bond-lifetime dynamics of tip-links that survive the periodic shearing forces at high-frequency and amplitude, and model their spring nature that can explain the uninterrupted hearing. The overarching objective of this work is to characterize tip-links as gating-spring and quantitatively evaluate their contribution over a wide range of sound intensity and frequency. The hypothesis is that the force sensed at the interface is likely the force at the MET channel. We, therefore, clamp the tip-links, a doublet of cadherin-couples, at constant tensile forces using atomic force microscopy (AFM) and monitored the temporal force-response of the binding interface. Simultaneously, we captured the force-induced alterations in the tip-link elasticity and correlated with the temporal response of the interface. The use of AFM is key to achieve the higher force-loading rates equivalent to high-frequency loud sound. In addition, we verified the effect of mutation associated with congenital hearing loss on the bond-lifetime dynamics. Moreover, the existing biophysical, genetic screening in model systems and other physiological studies verify that Cdh23 may be the dominant elastic component of tip-links(Arora et al., 2021; Astuto et al., 2002; Miyagawa et al., 2012). Here, we used different variants of Cdh23 to understand how alterations in the protein elasticity affect the bond-lifetime dynamics of tip-links.

## Results

### Force-induced transition from slip to catch bonds helps tip-links to survive sound

To distinctly measure the bond-survival dynamics of the tip-links complex, we first performed force-clamp experiments with cadherins containing two outermost domains (EC1-2). In particular, the two outermost domains of cadherins overlap with each other over a surface area of ∼1000Å^2^ and form the tip-links complex(Sotomayor et al., 2012). It is, thus, intriguing to deduce how a small binding interface of cadherins tolerates a wide range of force-stimuli from the sound of varying decibels. Accordingly, we attached Pcdh15 EC1-2 on the cantilever and Cdh23 EC1-2 on the surface, both covalently following a reported Sortase A mediated transpeptidation protocol(Srinivasan et al., 2017)and performed force-clamp measurements **(Figure 1c, d, Materials and Methods)**. We measured the force-tolerance time of the complex from each successful clamping and calculated the survival probability (SP) of the complex for each clamping force from repetitive measurements **(Figure 1e**). From the statistical F-test, we determined that the SP followed single-exponential decay. We subsequently measured the mean-lifetime of tip-links from the SP for varying clamping forces and obtained the numeric relation of force-lifetime of the complex. It is important to note that for every force-clamp experiment, we performed control experiments to quantify the non-specific contribution in the measurements **(Methods)**.

Lifetime of the complex follows a tri-phasic behavior with clamping force (**Figure 1f)**. Initially, the complex lifetime drops monotonically with increasing force, featuring slip bond. Slip bonding is consistent with the previous results obtained from the optical tweezer measurements at low-force regimes for slow force-loading rates(Mulhall et al., 2021). Above a critical force (*F*_*C1*_) of 52 pN, the bond-lifetime increases with force and reaches maxima at a critical force (*F*_*C2*_) of 90 pN, featuring catch-bond. A catch bond makes the interface tighter under the mechanical perturbation. Beyond 90 pN, the lifetime decreases monotonously with further force elevation, again slip behavior. So, the bond-lifetime of the complex does not monotonously slip with force, instead follows a roller-coaster ride of slip-catch-slip bonds. A slip to catch transition with force (>52 pN), thus, elevates the survival of tip-links especially at loud noise.

### Molecular details shed light on the slip-catch-slip transitions

What is the molecular mechanism of the slip-to-catch transition? To decipher, we performed Steered Molecular Dynamics simulations (SMD) using the infinite switch simulated twinning in force (FISST)(Hartmann et al., 2020) and quantified the effect of force on the tip-links interface. Similar to force-clamp experiments, we applied constant forces of 10 - 150 pN using the X component of the end-to-end distance as the collective variable (with the long axis of the complex aligned to the X axis). The complex was stable under these range of forces for microseconds, indicating no complex dissociation or domain-unfolding during simulation runs. Rather the external force tilts the energy landscape which is sampled by FISST simulations. We observed an increase in the buried surface area concomitant with an increasing tensile force (**Figure 2a**). Additionally, at intermediate forces of 40-90 pN, we observed another population of conformations with a distinctly larger buried surface area. An increase in buried surface area implies the strengthening of the complex in response to the applied force, in agreement with the experimental slip-to-catch transitions. Calculating the differential per residue buried surface area for the force range of 70-80 pN (compared to 10-20 pN), we noted that residues in both the EC1 and EC2 domains, as well as the N-termini of both proteins, are more buried upon the application of force (**Figure 2b**). Unlike experiments, we did not observe any SMD feature that explains the slip-behaviour of tip-links at low force regime. The absence of a slip phase in the MD simulations could be due to the formation of a weaker initial encounter complex in the force-clamp experiments which settles into a conformation similar to the crystal structure at very low forces. Such a transition is unexpected in our simulations as we start from the crystal structure.

**Figure 2:**
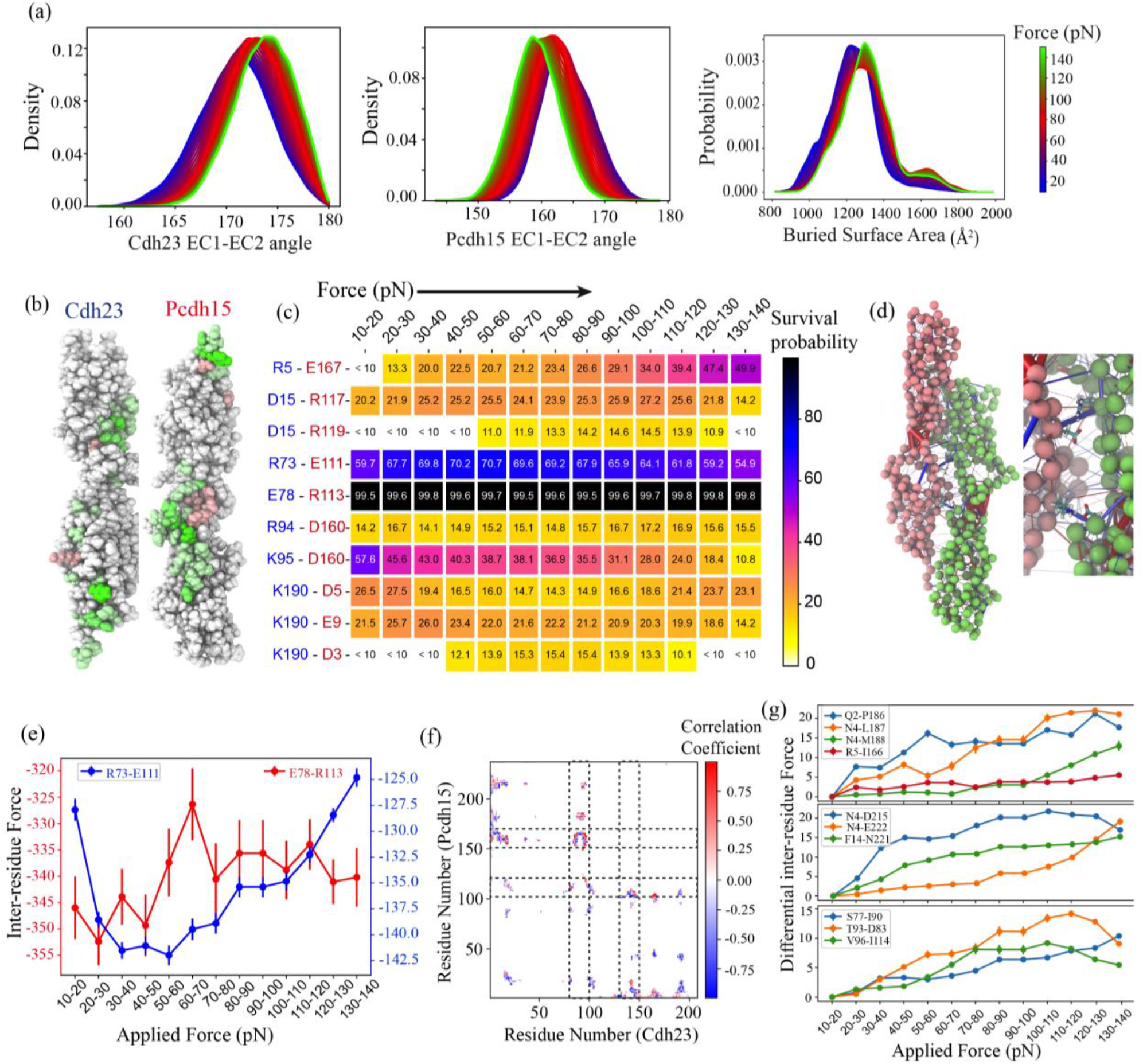
Load balancing in tip-links complex. **(a)** Distributions of inter-domain angles between EC1 and EC2 of Cdh23 (left) and Pcdh15 (middle) at the 61 interpolated points between the force range of 10 and 150 pN from FISST simulations. Variations in inter-domain angle is accompanied with the change in buried surface area (right). **(b)** The per-residue differential buried surface area calculated by subtracting the buried surface area at 10-20 pN force range from 70-80 pN force range. The residues with positive differential buried surface area are colored as green with darker shades representing higher values. The residues colored as red show a negative differential buried surface area with darker shades representing larger negative values. **(c)** The percentage of frames showing the existence of specified salt-bridges at different force ranges. **(d)** The pairwise inter-residue forces plotted over the structure of Cdh23-Pcdh15 complex. The color of the cylinders represents the sign of the force with blue being negative and red being positive. The width of the cylinders represents the magnitude of the force. **(e)** The inter-residue forces for the salt-bridge paired residues E78-R113 and R73-E111. **(f)** Correlation coefficient between the inter-residue forces 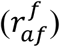 and the applied force for all the inter-protein residue pairs. **(g)** Selected residue pairs for which the inter-residue force is highly correlated with applied force 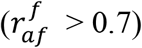. See also Figures S1-S3.

The molecular interactions in catch-regime are stabilized by tension. To identify, we calculated the probability of formation of salt-bridges (**Figure 2c**), hydrogen bonds **(Figure S1)**, and hydrophobic interactions **(Figure S2)** between Cdh23 and Pcdh15 at the force range of 10 - 140 pN in steps of 10 pN. Interestingly, a salt-bridge between Pcdh15-R113 and Cdh23-E78 remained intact at all forces emphasizing its importance in maintaining the complex even under tension. The probability of all the pre-existing salt-bridges increases slightly at small forces of 20-30 pN. At higher forces, some of the salt-bridges, viz. K95(Cdh23) - D160(Pcdh15), K190(Cdh23) - D5(Pcdh15), and K190(Cdh23) - E9(Pcdh15) became weaker with force (less probable). On the other hand, salt-bridges viz. R73(Cdh23)-E111(Pcdh15) and R5(Cdh23)-E167(Pcdh15) became stronger (more probable) with increasing force. Additionally, new salt-bridges appear at higher forces that are either absent or present in a negligible number of frames at low forces. This includes R5(Cdh23)-E167(Pcdh15), K190(Cdh23)-D3(Pcdh15), and D15(Cdh23)-R119(Pcdh15). Similarly, some of the hydrogen bonds between the EC1 domain of one partner and the EC2 domain of the other also became more probable at higher forces including V3(Cdh23)-P186(Pcdh15), F9(Cdh23)-Q218(Pcdh15) and Q188(Cdh23)-Y8(Pcdh15). A similar trend is seen for the hydrophobic interaction between V3(Cdh23)-L187(Pcdh15) **(Figures S1 and S2)**.

### Dynamic force balance protects essential interactions in the Cdh23-Pcdh15 complex

The salt-bridge between Pcdh15(R113) and Cdh23(E78) remains unaltered in the entire clamping forces. We propose the pivotal role of this salt-bridge on the slip-to-catch switch under tension. To understand the mechanics that conserve the pivotal role of this salt-bridge interaction at a large range of tension, we quantified how force is stored and transmitted in the complex. We estimated the pairwise inter-residue force distributions for each clamping force (**Figure 2d**). At the clamping force of 10-20 pN, which are extremely small forces for this complex, the two most important inter-protein salt-bridges i.e., Cdh23(R73) – Pcdh15(E111) and Cdh23(E78) – Pcdh15(R113) are most tensed i.e., they tend to pull the proteins closer. Interestingly, the salt-bridge between Cdh23(E78) – Pcdh15(R113) maintains a constant degree of tension across the entire loading force (**Figure 2e**). A bond bearing a constant tension across large tensile forces refers to a load balancing activity inside the protein complex that drives the force away from this critical interaction. On the other hand, the force between Cdh23(R73) – Pcdh15(E111) gradually becomes less negative with the applied force, indicating that this interaction will break at higher forces or at a longer time (**Figure 2e**). In order to decipher the inter-residue interactions that might absorb the applied force and steer it away from these critical salt-bridges, especially from E78-R113, we calculated the correlation of pairwise inter-residue forces 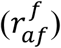 with the applied force (**Figure 2f)**. The residue pairs with 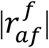 significantly greater than zero, experience a change in the inter-residue forces as a function of applied force. These residue pairs should be responsible for actively distributing the load inside the proteins and the complex, thus assisting Cdh23(E78) – Pcdh15(R113) to remain unperturbed. On the other hand, residue pairs with 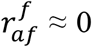 bear a consistent load irrespective of the applied force. Using the scale of 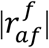, we, therefore mapped the correlations among the load balancing residues (**Figure 2f)** and identified crucial interactions between the EC1-EC1 and EC1-EC2 domains of Cdh23 and Pcdh15, respectively. Additionally, to highlight the residue-pairs that are strongly correlated to the applied force, we mapped the differential inter-residue force (inter-residue force at a force range – inter-residue force at 10-20 pN) (**Figure 2g)** and observed a strongly interconnected network in certain regions of the proteins (e.g., residues 150-175 and 100-125 for Pcdh15 and 75-100 and 130-150 for Cdh23) (black dashed lines, **Fig 2f**). This network of interconnected residue pairs bears and distributes the applied force to maintain the integrity of the complex. For reference, we also mapped the correlation among load bearing residues within the protein **(Figure S3)**.

### The slip-to-catch transition is abolished in an inherited deafness mutation in Pcdh15 (R113G)

In reference to the SMD results, inter-protein salt-bridge Cdh23(E78) – Pcdh15(R113) remains intact with a dominant survival throughout the force range. The proteins undergo inter-domain rotations to make new contacts or strengthen existing ones to allow the complex to resist the applied force. From these observations, it is apparent that this robust bridge serves as a molecular fulcrum which drives the switch from a weaker binding-interface to a stronger binding-interface of the complex under the tensile force. A transit from a weaker binding interface to a stronger interface is responsible for the catch-bond behavior. The tensile force here acts as a catalyst and exposes the cryptic binding interface by imparting entropic energy for extension and rotation of the interacting partners. Incidentally, the importance of R113 of Pcdh15 is also known in physiology, where the mutation of R113 to glycine (R113G) is prevalently found in patients suffering from inherited deafness(Ahmed et al., 2008). Accordingly, we recombinantly synthesized Pcdh15 EC1-2 (R113G) and studied the bond-lifetime dynamics of the mutant tip-links Cdh23 EC1-2 (WT)-Pcdh15 EC1-2 (R113G) with force using force-clamp spectroscopy. Pcdh15 R113G fails to steer the slip-to-catch transition with force and the complex monotonously slips towards dissociation with force **(Figure 1f**). It is, therefore, justified to propose that the slip-catch-slip feature of the wild-type tip-links is essential for normal hearing and once the slip-to-catch transition is lost, the hearing is acutely compromised.

### A kinetic (force-induced stronger-binding) model to explain the slip-catch-slip transitions

The kinetic models of catch, slip, and ideal bonds were proposed prior to even any experimental observation. A dynamic switch in the mechanical stiffness of the biological bonds under tension was proposed as responsible for all three types of bonds(Dembo et al., 1988). Subsequently, several phenomenological theories, including a two-pathway model(Pereverzev et al., 2005), allosteric bond model(Thomas et al., 2006), deformation-model(Nair et al., 2016), sliding-rebinding model(Lou and Zhu, 2007) were derived to explain the catch bonds. From in-silico studies, we learnt that the force-induced transition to a denser binding interface is responsible for catch. Accordingly, we derived a ‘force-induced stronger-binding model’ from the ‘sliding-rebinding model’ and fit our data **(Methods)**. From barrier-crossing perspective, the complex may follow the two pathways. Pathway one involves the force-induced conformational transition to a denser binding interface. The pathway two is the direct dissociation. We define a critical force (F_C1_) to tune the probabilities of denser-binding or direct dissociation, and a second critical force, F_C2_, to limit the catch regime (**Eq 1, Methods)**. Above F_C2_, bond-lifetime decreases monotonously. The rates of sliding (k_+1_) and rebinding (k_+2_) are considered independent of force whereas the rates of dissociation (k_-1,_ k_-2_) in both the pathways are force-dependent **(Methods)**. Our model fits our experimental data convincingly (**Figure 1f and Table 1**). We fitted the slip-bond lifetime-force data of the mutant complex with Bell’s model(Bell, 1978) and obtained the intrinsic off-rate, 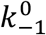 and distance to transition, *x*_*β*_ (**Figure 1f**). Interestingly, 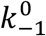 (0.3 s^-1^) and *x*_*β*_ (0.8 Å) of mutant tip-links are comparable to that of WT tip-links (0.2 s^-1^ & 1.1 Å, respectively) **(Table 1**). However, the bond-survival for mutant drops rapidly with force while WT tip-links survive high-intensity stimuli by switching to catch conformation under tension. Rapid unbinding of mutant tip-links under high-intensity stimuli is also measured previously(Mulhall et al., 2021). It was, therefore, inferred that the severity of the homo-/heterozygous Pcdh15 R113G mutant is more prominent in cochlear hair-cells where the exertion of force is larger than the vestibular hair cells(Sahly et al., 2012).

**Table 1:** Kinetic parameters obtained from the force-induced rebinding fit of lifetime-force data for Cdh23 EC1-2-Pcdh15 EC1-2.

| $k_{-1}^0 (s^{-1})$ | $x_\beta (\text{\AA})$ | $k_{+1} (s^{-1})$ | $k_{+2} (s^{-1})$ | $F_{c2} (pN)$ | $n$ |
| --- | --- | --- | --- | --- | --- |
| 0.2 | 1.1 | 0.1 | 10 | 116 | 2.1 |

### Unfolding mediated unbinding controls the force-dissemination process in tip-links

Dynamic alterations in the entropic conformations of the extraordinarily long tip-links cadherins under tension are already reported(Arora et al., 2021; Bartsch et al., 2018; Choudhary et al., 2020; Oroz et al., 2019). How such alterations contribute to the bond-lifetime dynamics of tip-links is not yet clear. Towards this, we systematically altered the number of the non-interacting domains of Cdh23 and probed the tip-links survival using force-clamp spectroscopy. Apparently, Cdh23 possesses comparatively larger entropic conformations and least axial stiffness among tip-links proteins. We recombinantly synthesized four different variants of Cdh23, Cdh23 EC1-5, Cdh23 EC1-10, Cdh23 EC1-21, and full-length Cdh23 EC1-27 **(Methods)**. It is apparent that there will be a monotonous drop in protein-stiffness of Cdh23 with increasing number of domains and linkers. For force-clamp experiments, we used the two outermost domains of Pcdh15 (EC1-2) on cantilever as pulling handle and Cdh23-variants independently on coverslip **(Figure 3a, Methods)**. The slip-catch-slip features persisted for all the complex-variants **(Figure 3d)**, indicating the bond-behaviors as trademark for the interface of individual tip-links. However, overall lifetime of the complex is systematically elevated with increasing number of domains **(Figure 3d)**.

**Figure 3:**
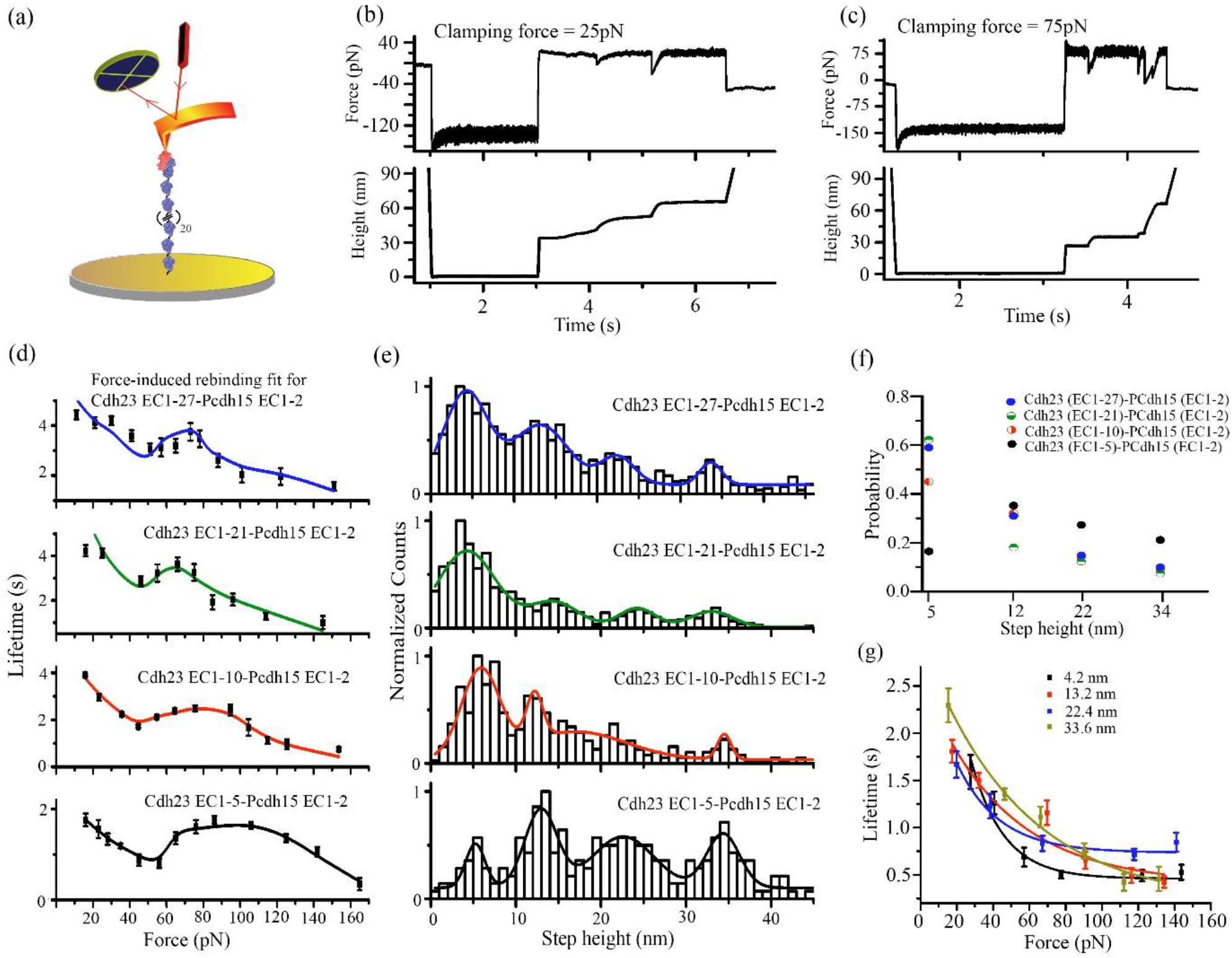
Effect of molecular-elasticity on the tip-links interface. **(a)** Schematic representation of AFM-based force-clamp experiment for Cdh23 EC1-27-Pcdh15 EC1-2 complex. We performed the force-clamp measurements with Cdh23 constructs of varying domain numbers (n) of 5, 10, 21, and 27. **(b and c)** Representative force vs. time and height vs. time plot for clamp measurements of Cdh23 EC1-27-Pcdh15 EC1-2 interaction complex. Unlike the previous force-clamp measurements with only two EC domains, here we observe instantaneous drop of force followed by restoration in the force vs time plot. Similar feature is also reflected in the corresponding length vs. time plot with stepwise height change. Such features indicate extension of protein during the clamping condition. **(d)** The lifetime-force behavior for tip-links complexes for different lengths of Cdh23 constructs. All the complexes depict slip-catch-slip transition with increasing force. Error bars represent the standard errors obtained from the exponential fitting of the survival plots. **(e)** Unfolding step-height distributions, calculated from the force curves for all Cdh23 variants, Cdh23 EC1-5 (black), Cdh23 EC1-10 (red), Cdh23 EC1-21 (green), and Cdh23 EC1-27 (blue). Solid lines are the Gaussian fits to the distribution providing the most probable step-height from the peak maxima. **(f)** Relative probabilities of each type of step-height change for clamp measurements with varying lengths of Cdh23 construct. **(g)** Force-lifetime data for all four major extensions from Cdh23 EC1-27-Pcdh15 EC1-2 force clamp experiment to obtain the kinetic parameters like unfolding rate at zero force 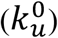 and distance to transition state (*x*_*β*_). See also Figure S4 and S5.

We observed four extensions of tip-links proteins of ∼5 nm, ∼12 nm, ∼22 nm, and ∼ 34 nm (**Figure 3b,c,e and Table 2**) for all complex-variants, prior to dissociation. Extensions increase the complex lifetime (**Figure S4)**. Although the probabilities of the respective extensions varied across complex-variants, the probability of extension is maximum for ∼5 nm and insignificant for ∼22 nm and ∼34 nm for all variants (**Figure 3f**). To accommodate the unfolding kinetics in the tip-links survival, we modified the force-induced stronger-binding model by including the experimental extension probabilities **(methods)**. We segregated the lifetime-force data for all the step-heights ∼5, ∼12, ∼22, ∼34 nm for full-length Cdh23 EC1-27 and fitted to exponential decay (Bell model) to obtain the zero-force unfolding rate 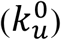 and distance to transition-state (*x*_*u*_) for the individual step height (**Figure 3g, Table 3)**. For simplicity, we included the two most-probable extensions of ∼5 nm and ∼12 nm in the model and fit to the force-induced bond-dissociation dynamics of all the complex variants, independently (**Figure 3d, see Methods**). From the fits, we obtained the intrinsic lifetimes of the tip-links interface which increases with increase in the number of EC domains. The zero-force lifetime is increased from ∼1.5 s for Cdh23 EC1-5 to ∼10 s for Cdh23 EC1-27 variant. The critical force (F_C2_) at which the catch switches to slip bonds shifts lower, 108 pN for Cdh23EC1-5 to 68 pN for Cdh23 EC1-27. Further, we noted a decreasing trend in the distance to transition state (*x*_*β*_) and rebinding rate (*k*_+2_) with increasing domains **(Table 4)**. On contrary, the parameter referring to the force-enhanced binding, *n*, increases with the domain number, with the smallest value of 1.0 for 5 EC domains to the highest value of 5.8 for 27 EC domains. The increment in *n* justifies the enhanced force-resistance of tip-links towards dissociation (higher bond-lifetime). The lowest *x*_*β*_, 0.5 Å, for the full-length variant too signifies the importance of the extended EC domains of tip-links proteins in force-tolerance. The sliding rate (*k*_+1_) or the re-organization rate of the binding interface, however, remains unaffected with increasing domains. This was intuitive from the SMD as the inter-protein pivot, the salt-bridge between Cdh23 (E78)-Pcdh15 (R113), did not show any alteration with increasing applied forces. Further, we noticed a gradual increase in the percentage of unfolding associated unbinding events with increasing EC domains **(Figure S5)**. Increasing domains with linkers systematically lower the persistence length (*l*_*p*_) of the protein variants, and so the tension propagation time(Bohbot-Raviv et al., 2004), 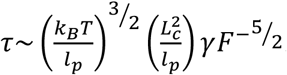. A delay in tension propagation to the binding interface eventually slows down the dissociation. Moreover, the extensions did function as dissipators of force and uplift the bond-survival significantly under moderate tension **(Figure 3d)**. However, at extreme stimuli, tip-links delink faster.

**Table 2:** Most probable step heights from peak maxima of the unfolding step height distributions for different tip-links complexes.

|  | Step height 1<br>(nm) | Step height 2<br>(nm) | Step height 3<br>(nm) | Step height 4<br>(nm) |
| --- | --- | --- | --- | --- |
| Cdh23 EC1-5 | 5.2±0.3 | 12.9±0.3 | 22.6±0.5 | 34.3±0.3 |
| Cdh23 EC1-10 | 5.8±0.3 | 12.2±0.2 | 17.1±2.9 | 34.4±0.9 |
| Cdh23 EC1-21 | 3.7±0.2 | 10.0±0.6 | 24.6±0.7 | 32.8±0.8 |
| Cdh23 EC1-27 | 4.2±0.2 | 13.2±0.4 | 22.4±0.4 | 33.6±0.4 |

**Table 3:**
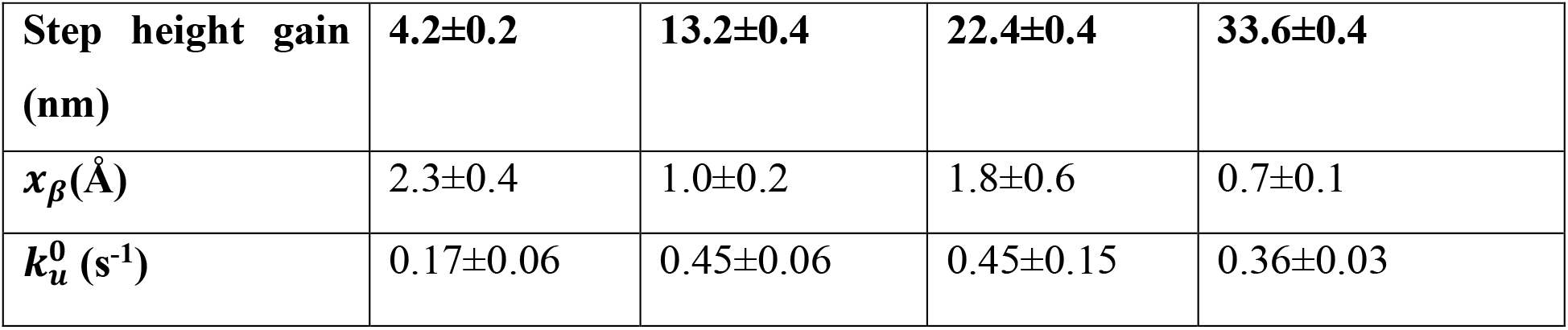
Kinetic parameters obtained from Bell’s-model fitting of force-dependent lifetime data of individual step height in Cdh23 EC1-27-Pcdh15 EC1-2 complex.

**Table 4:** Kinetic parameters obtained from the modified force-induced rebinding fit of lifetime-force data for different variants of Cdh23 in complexation with Pcdh15 EC1-2.

| | $k_{-1}^0 (s^{-1})$ | $x_\beta (\text{\AA})$ | $k_{+1} (s^{-1})$ | $k_{+2} (s^{-1})$ | $F_{c2} (pN)$ | $n$ |
| --- | --- | --- | --- | --- | --- | --- |
| Cdh23 EC1-5 | 0.6 | 1.2 | 1.2 | 20 | 108 | 1.0 |
| Cdh23 EC1-10 | 0.4 | 1.0 | 1.0 | 7.8 | 99 | 1.9 |
| Cdh23 EC1-21 | 0.2 | 0.8 | 1.1 | 1.0 | 69 | 3.8 |
| Cdh23 EC1-27 | 0.1 | 0.5 | 0.7 | 0.2 | 68 | 5.8 |

### Heterotetrameric tip-links are insensitive to force

Tip-links exist as heterotetramer in inner ear(Kazmierczak et al., 2007). A cis-homodimer of Pcdh15 forms a fork-like structure with their two outermost EC domains (EC1-2) and facilitate the trans-binding with two opposing Cdh23 proteins(Kachar et al., 2000; Sotomayor et al., 2012). Such a tetrameric conformation of tip-links confines two protein-complexes spatially, and elevates the complex-lifetime through stochastic rebinding(Mulhall et al., 2021). However, the effect of stochastic rebinding on the force-induced bond-dissociation dynamics is still speculative. To attempt, we replicated the tip-links geometry in a force-clamp setup, i.e., the cis-dimers of Cdh23 EC1-27 at cantilever and Pcdh15 EC1-11 PICA dimers on coverslip **(Methods, Figure 4a)**. For cis-dimer of Cdh23, we recombinantly tagged the C-terminal of the protein with the fragment crystallizable region (Fc-region) of an antibody. The Fc constructs self-ligate via disulfide linkages and brings a pair of Cdh23 proteins in close proximity **(Methods, Figure S6)**. Pcdh15 EC1-11 PICA predominantly exists as cis-dimer in the ambient laboratory conditions (**Figure S6**). Tip-links doublets follow slip bonds at low-force regime **(Figure 4c)**. At the intermediate forces (∼36 pN to ∼70 pN), the stimuli we regularly perceive, the doublet complex features an ideal-bond character, an indifferent force-response by the complex interface **(Figure 4c)**. While the molecular mechanism is not clear yet, we speculate that the ideal bond feature is the manifestation of force-filtering. Tip-links could diffuse the tensile forces dynamically and efficiently so as to expose the binding-interface to a constant force in the ideal-bond regime, an action similar to low-force pass filter. At extreme auditory inputs, beyond 100 pN, tip-links progressively dissociate with force, characteristic to slip bond.

**Figure 4:**
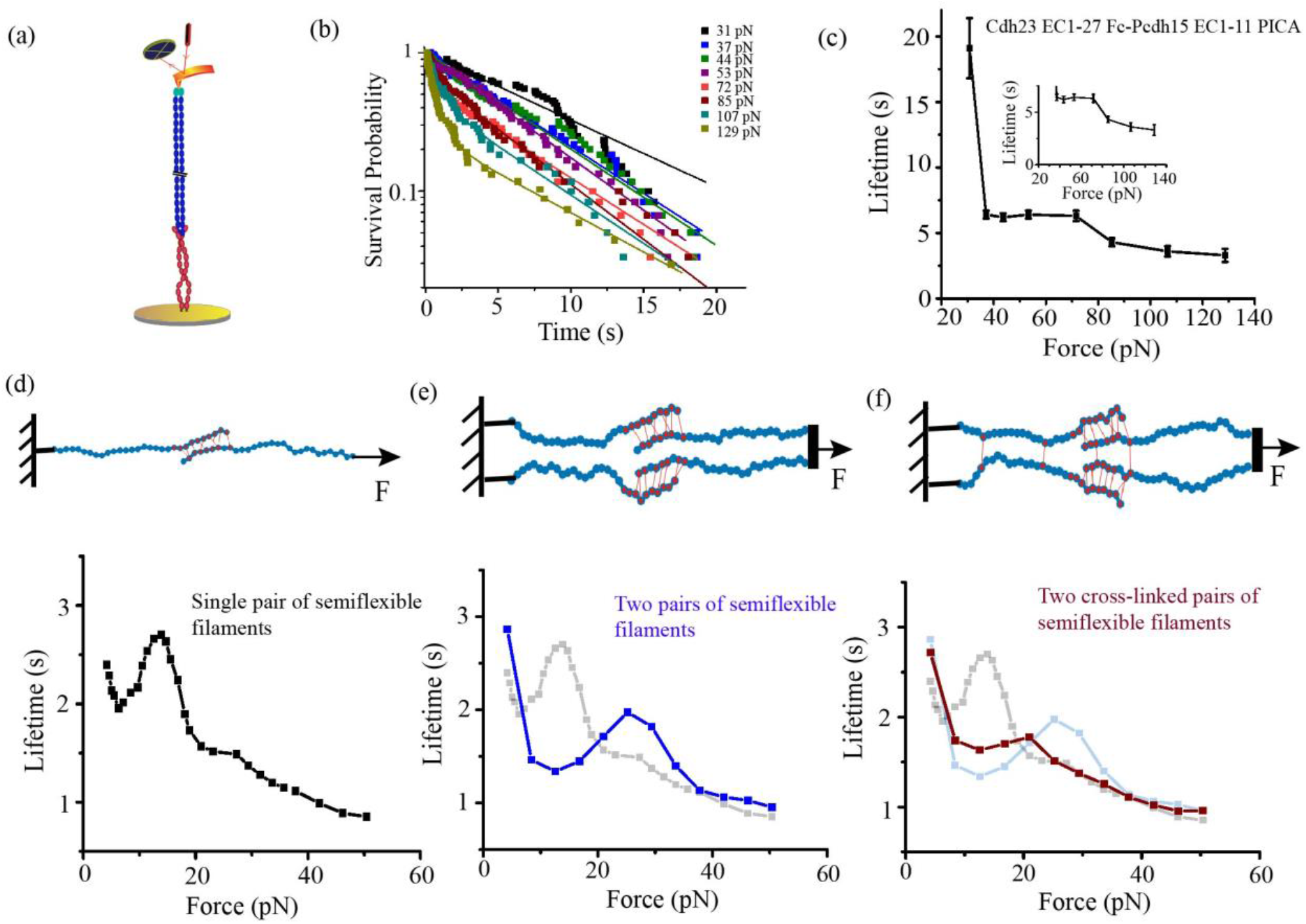
Heterotetrameric tip-links complex features a slip-ideal-slip bond. **(a)** Schematic representation of the tetrameric tip-links complex. Cdh23 EC1-27 (blue) with Fc region (green) on its C-terminus is covalently attached to the cantilever while the dimer of Pcdh15 EC1-11 (PICA) is attached on the surface. **(b)** The bond-survival probability plot at eight clamping forces fitted with exponential-decay model to obtain the bond lifetime. The solid lines represent the exponential-decay fit. **(c)** The tetrameric assembly of tip-links forms slip-ideal-slip bonds with increasing force. Inset shows the zoomed view of the ideal bond regime. Error bars represent the standard errors obtained from the exponential fitting of the survival probability. **(d)** Schematic representation (top) of two semiflexible filaments coupled with multiple elastic slip-catch bonds resembling the individual tip-links, and clamped at constant forces to perform the coarse-grained Langevin dynamics simulations. Bottom curve shows the resulting force-dependent lifetime that depicts the slip-catch-slip behavior. **(e)** Schematically representing the two pairs of the semiflexible polymers coupled and pulled in combination at constant forces. Slip-catch-slip feature remain unaltered (blue curve, bottom) for this polymer system, however, lifetime in the catch-regime is dropped compared to single pair of filaments (grey curve in background). **(f)** A modified polymer model from (e) where the chains from the same sides are connected. This model mimics the heterotetrameric geometry of the tip-links. The characteristic force-lifetime curve exhibits a slip-ideal-slip feature (dark red, bottom). Grey and light blue curves in the background are from model (d) and (e). See also Figures S6-S10.

We observed extensions, as previously, ∼ 5 nm as most preferred followed by ∼12 nm extensions. Other extensions as seen in monomeric tip-links are vanishingly small for heterotetramer complex **(Figure S8)**. Interestingly, the ∼5 nm extension, referring to inter-domain linker elongation, is reported as elastic stretching(Bartsch et al., 2018). Our force-distribution analysis too identified the linkers as pre-compressed between EC1-2 domains of both the proteins **(Figure 2d)**, marking linkers as elastic springs in force-dissipation. However, this pre-compression is not only limited to two domains, rather when we go away from the interface i.e. linker between EC2-3 domains, there also we observed that the linker is pre-compressed **(Figure S9a)**. Further, the ∼12 nm extension, referring as partial domain unfolding(Arora et al., 2021), may diffuse force non-elastically and could be the dashpots in the spring. Notably, a gating-spring by definition, must possess elastic and non-elastic (dashpot) compliances. We may therefore, picturized the tip-links in spring-dashpot model where the tip-links as ‘gating-springs’ are connected with multiple elastic springs (Δx ∼5 nm) and relatively a smaller number of dashpots ((Δx ∼12 nm) in a series **(Figure S9b)**.

The extrapolation of slip bonds at low-force regime results a long-lasting intrinsic lifetime (τ_0_ ∼ 30 s) for the tetrameric tip-links, at no force. However, lifetime estimated from the curve may still be under-valued as almost one-third of the clamps did not dissociate within the clamp-time of 20 s in the low-force regime. Further, we noted that survival probability of tip-links complexes at high forces (> 70 pN) followed bi-exponential decays with time, instead of mono-exponents as in low-force region **(Figure 4b)**. We estimated a faster dissociation is contributing towards the second component. Intuitively, the short-lived component is arising from the dissociations of tip-links without re-binding. This is also reflected in the corresponding amplitude which increases with the force **(Figure S7)**. Lesser rebinding probability at higher tensile forces were noticed previously from in-silico kinetic models(Mulhall et al., 2021).

### Tetrameric assembly of tip-links crucial in gating-spring activity: A coarse-grained study

What is the origin of the ideal bonds? We already observed in the previous section that the molecular-elasticity diffuses the applied tension to uplift the survival, however, hardly alters the force-response nature of the binding interface. It is thus logical to hypothesize that the switch from slip-catch to slip-ideal bonds may be steered through dimerization of two independent tip-links. To understand the emergence of the slip-ideal-slip behavior of the tetrameric assembly of tip-links, we perform coarse grained Langevin dynamics simulations of coupled polymeric systems in two dimensions. We first consider an arrangement of two semiflexible filaments partially attached to one another via elastic bonds with catch-slip dissociation characteristics **(see Methods, Figure 4d)**. The bonds dissociate in the presence of elastic load when one of the filaments is pulled externally. The average life time of the attachment defined as the time beyond which all bonds between the two filaments break leading to two separated filaments, is plotted as a function of the external load force. The lifetime decreases initially with increasing force. Beyond a force, the lifetime increases showing the onset of catch behavior **(Figure 4d)**. As the external force is increased further, the rate of detachment starts increasing leading to lower survival time at large forces. Such an arrangement mimics individual tip-links at a coarse-grained level. It is important to note that the characteristic slip-catch-slip feature is independent of the stiffness of the individual filaments.

After training our dimeric system of two coupled filaments under external force, we next consider the tetrameric arrangement as shown in **Figure 4e**. Here, two filaments from each of the dimeric arrangement are pulled at one end simultaneously with the external force. We first look at the situation where there are no elastic bonds cross-linking the two dimeric systems, except for the end monomers which are held at the same position for both. The force-response of the bonded interface still follow a slip-catch-slip characteristic, although this feature is significantly reduced in the tetrameric arrangement with the peak in the lifetime shifting to higher force value. Finally, we cross-linked the polymers that are anchored, near terminals but away from cross-coupling region, resembling cis-dimers of Pcdh15 **(Figure 4f)**. We also cross-linked the polymer couples that are pulled at their cross-linking interface. This polymer couple resembles Cdh23. Cross-linking the dimeric systems via elastic bonds leads to an even further reduction in the catch-slip peak. For a choice of parameter values, we noticed the emergence of slip-ideal-slip behavior as observed in the experiments.

## Discussion

Force-response of a binding-interface is a direct measure of the force-modulation by the viscoelastic force-transducer. A slip-ideal bond interface of heterotetrameric tip-links reveals the low-force pass filter like feature for tip-links. A low-force pass filter exposes a range of low forces to the binding-interface where the interface slips with force. In the ideal-region, tip-links dynamically dissipate forces and exert a constant force to the interface. A low-force pass filter thus aptly demonstrates the working mechanism of a gating-spring, which dissipate the surplus of tension from loud noise while conveying a threshold to MET channels, and without leaving any adverse effect on cell-membrane. The ideal bond interface of tip-links thus ensures steady hearing across a range of auditory inputs. However, at very loud sound, tip-links follow slip bonds and progressively dissociate to protect any permanent damage in the sensory organs. The binding interface of the individual tip-links complex forms slip-catch bonds with increasing force, a mechanism to delay dissociations in moderate to loud sound. The catch bond is essential to achieve the ideal bond feature in the tetrameric tip-links (**Figure S10**). Using all-atom non-equilibrium simulations, we revealed that the slip-to-catch switch under external force is modulated by a ‘molecular fulcrum’, a salt-bridge between Cdh23(E78) and Pcdh15(R113). Interestingly, mutation of R113 to Gly is linked to hereditary deafness. We, too, noticed the obsolete of catch regime with Pcdh(R113G), emphasizing further the physiological importance of the catch bond-interface.

Unlike catch bonds or slip bonds, ideal bonds are least explored by nature. In literature, ideal bonds are generally referred as experimental artifacts where the direction of force-application and bond-dissociation are orthogonal to each other(Dembo et al., 1988; Rakshit and Sivasankar, 2014; Rakshit et al., 2012). However, the origin of the ideal bond in tip-links is not an artifact, it is rather unique. Ideal bond emerges from the lateral dimerization of two individual tip-links complexes, however, only when the interprotein couplings between two Pcdh15 and two Cdh23 were established. The ideal bond is thus, a cooperative effort of two individual tip-links complexes, laterally adjoined by nature. For Pcdh15, these interprotein lateral couplings are at the PICA domain and EC3 domain, away from the interacting interface. For Cdh23, such lateral coupling is formed between EC1 domain, however, only when in close-proximity. It is, thus, apparent that the cis-dimer cross-links are key to diffuse the applied forces orthogonally along cross-links(Arora et al., 2021; Hazra et al., 2019; Ott et al., 2015). We, therefore, propose that the heterotetrameric geometry of tip-links is designed and fabricated by nature on a purpose to serve as simplest gating-springs.

## Conclusion

We deciphered the functional motive of the unique configuration of cadherin duos in tip-links. Tip-links as gating-spring behaves like a low-force pass filter. Our single-molecule force-clamp experiments and detailed simulations clearly demonstrated that the force-insensitive ideal-bond feature is an aftermath of the hetero-tetrameric assembly of cadherins with significant contributions from the homotypic cross-links among cis-dimers. However, the overarching mechanism of such cooperative force-transmission is not clear yet.

## Methods

### Cloning, expression and purification of Cdh23 EC1-2 and Pcdh15 EC1-2 using bacterial system

Mouse Cdh23 EC1-2 and Pcdh15 EC1-2 were recombinantly modified with 6xHis-tag at N-terminal for affinity-purification, and sort-tag (LPETG) at C-terminus for surface attachment and cloned into Nde1 and Xho1 sites of pET21a vector. Mutation R113G in Pcdh15 EC1-2 was engineered using overlapping extension PCR. All the constructs were verified by DNA sequencing. Proteins were expressed in *E*.*coli*. BL21 Codon Plus (DE3)-RIPL cells (Stratagene, San Diego, CA, USA) cultured in LB media. Cells were induced with 200 µM IPTG at OD 0.6 at 16°C for 16 h. Cells were lysed by sonication in the 8M urea buffer and protein was purified from inclusion bodies using Ni-NTA based affinity purification. Eluted proteins were refolded by sequential dialysis in decreasing concentration of urea and further purified with size exclusion chromatography. The purity of the proteins was monitored from SDS-PAGE gel electrophoresis.

### Cloning, expression and purification of Cdh23 truncated variants in mammalian system

For Cdh23 EC1-27 construct, we ligated the two fragments of Cdh23 having overlapping regions using Gibson assembly(Gibson et al., 2009). From the Cdh23 EC1-27, we generated all the truncated variants; EC1-5, EC1-10, and EC1-21 between restrictions sites NheI and XhoI in vector pcDNA3.1(+) using the DNA recombination method. All the constructs have sort-tag (LPETGSS), GFP and 6xHis-tag at the C-terminus. Sort-tag was inserted for covalent attachment to surface, GFP tag helped in tracking the protein expression and His-tag was necessary for affinity-based protein purification. All the constructs have signal peptide at the N-terminus to guide the expressed protein in the media.

For the heterotetrameric tip-links construct, we recombinantly modified the C-terminus of Cdh23 EC1-27 with fragment crystallizable region (Fc-region) of a human antibody. The Fc constructs can self-ligate via disulfide linkages and bring a pair of Cdh23 proteins in close proximity. For Pcdh15 EC1-11 PICA, we PCR amplified the full Pcdh15 plasmid up to the PICA domain using specific primers and cloned in pcDNA 3.1 (+) vector between the restriction sites KpnI and XhoI.

For protein expression, we used Expi-CHO suspension cells (Thermo Fisher Scientific). Cells were transfected at the cell density of 10^8^ with the 1µg/mL plasmid as the prescribed protocol and kept at 37°C, 6% CO_2_, 125 RPM in the incubator. After 20 h of transfection, feed and enhancer was added to increase the yield of protein expression. Media was collected after 7 days from the transfection and centrifuged at 2000 RPM for 10 min. After centrifugation, we dialyzed the supernatant media in the buffer (25 mM HEPES, 150 mM NaCl, 50 mM KCl, 5 mM CaCl_2_) at 4°C. Dialyzed media carrying the protein of interest was purified using Ni-NTA based affinity purification.

### Surface modification for single-molecule force spectroscopy using AFM

Glass coverslips were activated using air plasma and subsequently washed with piranha solution for 2 h, followed by a thorough wash with deionized water. Coverslips were then etched using 1 M KOH for 15 min and washed with deionized water by sonicating for 10 minutes three times. Subsequently, the surfaces were silanized using v/v 2% APTES (3-Aminopropyltriethoxy silane) in 95% acetone and cured at 110 °C for 1 h. The amine exposed surfaces were reacted with 10% Maleimide-PEG-Succinimidyl ester (Mal-PEG5000-NHS) in methoxy-PEG-Succinimidyl ester (mPEG5000-NHS) (LaysonBio) in a basic buffer (100 mM NaHCO_3_, 600 mM K_2_SO_4_, pH 8.5) for 4 h for the experiment with just two interacting domains. For experiments with the varying lengths of Cdh23, we have used Mal-PEG2-NHS (Sigma-Aldrich) instead of (Mal-PEG5000-NHS). The PEGylated surfaces were subsequently incubated with 100 µM polyglycine peptide, GGGGC, at room temperature (RT) for 7 h for cysteine – maleimide reaction. Polyglycine on coverslip acts as a nucleophile for sortase-based covalent attachment. Coverslips were then washed with water and stored in a vacuum desiccator prior to protein attachment.

We used less stiff Si_3_N_4_ cantilevers (NITRA-TALL from AppNano Inc., USA) for all the force-spectroscopy studies. Cantilevers were functionalized in the similar way with GGGGC to attach Pcdh15 EC1-2 covalently using sortagging.

### AFM based force–clamp spectroscopy

C-terminus of Cdh23 was covalently immobilized on the freshly prepared glass coverslip coated with polyglycine and Pcdh15 was decorated on cantilevers using sortagging protocol in presence of enzyme sortase A as described previously(Srinivasan et al., 2017). Force-clamp spectroscopy was performed using Atomic Force Microscope (AFM) (Nano wizard 3, JPK Instruments, Germany). Spring constant of the cantilever was determined using equipartition based thermal fluctuation method(Hutter and Bechhoefer, 1993). After modifying the surface and cantilever with the proteins of interest, cantilever was approached towards the surface to interact with its partner. We let the proteins interact for 1-2 s, thereafter, retracted the cantilever by 20-70 nm depending on the proteins attached to cantilever, and clamped at a certain force for 10 s until the bond dissociation occurred, and then finally retracted the cantilever to break all the remaining bonds formed. For experiments with Cdh23 EC1-27 Fc dimer, we retracted the cantilever 70 nm up prior to clamping and set a clamping time of 20 s. All the experiments have been performed in the buffer with 25 mM HEPES, 100 mM NaCl, 50 mM KCl and 50 µM CaCl_2_. We repeated the experiment at different points on the surface and recorded around 2,000 force-curves at each clamping force to get statistically significant data.

For the control experiment, force-clamp data was recorded in without calcium buffer, 25 mM HEPES, 100 mM NaCl, 50 mM KCl along with 1 mM EGTA. We found that in absence of calcium, event rate decreased to ∼0.5-1% from 6-8% in the presence of calcium. This indicates that our measurements are specific to the interaction of tip-links cadherins.

After data acquisition, we measured the lifetime of the bond as the difference between the time when clamping force reached and when spontaneously bond dissociates. This persistence time of the bond gives the bond lifetime. We pooled all the data and divided in force bin. Thereafter, we plotted the survival plot at each force bin as the number of intact bonds vs. time. Exponential fit of the survival probability gives the bond lifetime at a certain clamping force. We have done F-test to estimate the best model of the exponential decay and found that for all the experiments with monomeric tip-links, mono-exponential decay is the best fit as compared the goodness-of-fit between the mono and bi-exponential decay. While for heterotetrameric tip-links, mono-exponential decay best fits the survival-time data at low forces while it fits with the bi-exponential decay at higher forces.

### Force-enhanced stronger-binding Model

For unbinding kinetics of tip-links with two-interacting domains, we modified the already existing sliding-rebinding model(Lou and Zhu, 2007) to fit our triphasic-bond transition data by introducing two critical forces, one corresponds to the minimum threshold force for slip to catch transition (F_C1_) while second is the force after which second slip started to appear (F_C2_). When force is less than F_C1_, no slip to catch transition would occur and direct dissociation pathway is followed while above F_C1_, unbinding occur through an alternate pathway where sliding or conformational rearrangement is followed by enhanced-binding that result in the observed catch behavior.

We consider the n pairs of pseudoatoms to describe the noncovalent interactions between the two protomers to fit the slip-catch-slip lifetime-force data. This will correspond to the *S*_*n*_ state in the model schematic. With the force application, m bonds break and the complex transitions to the *S*_*n*−*m*_ state. Further, from this state, there are two possibilities, either the complex undergoes to a completely dissociated state (*S*_00_) or to a stronger-binding state (*S*_*n*′_) through a force-induced conformational rearrangement (*S*_*n*−*l*_).

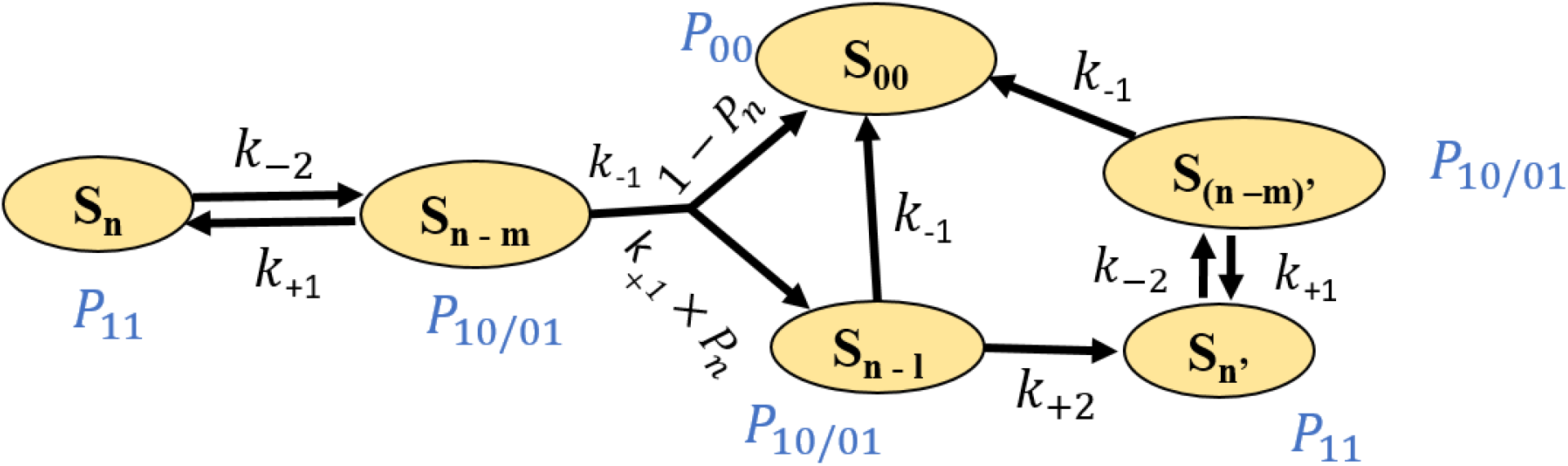

Kinetic rate equations for probability of different states are:

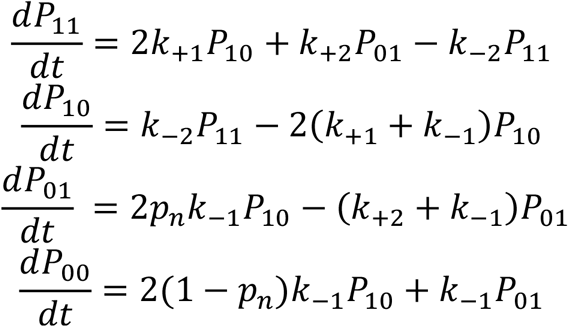

*k*_+1_ and *k*_+2_ is constant association rate and *k*_−1_ is force-dependent dissociation rate that followed the Bell equation(Bell, 1978):

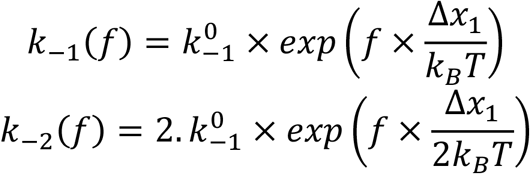

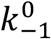 is the zero force off-rate and Δ*x*_1_ is the distance from bound state to the transition state. Probability (P_n_) of making new bonds and undergoing stronger-binding is defined as following:

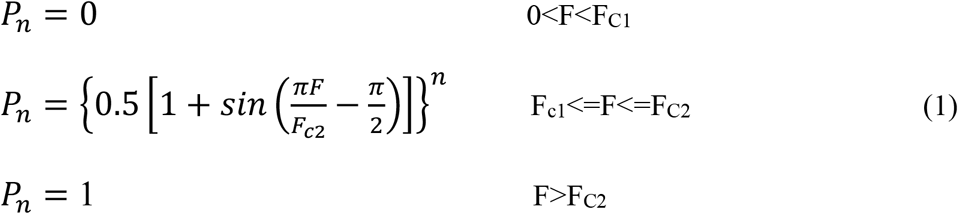

Here, n is the fitting parameter and Fc1, F_C2_ are the critical forces that define the first slip to catch transition and catch to second slip transition respectively. Probability rate equations were solved analytically using MATLAB and survival probability was estimated as

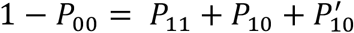

### For unfolding associated unbinding kinetics of tip-links involving non-interacting domains

We calculated the force-dependent unfolding probabilities for the most predominant unfolding step heights (∼5 and ∼12 nm) in accordance with the following equations(Brujić et al., 2007; Cao et al., 2008):

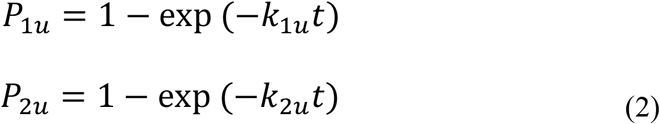

Where *P*_1*u*_, and *P*_2*u*_ are the unfolding probabilities of 5 nm and 12 nm step height respectively. *k*_1*u*_ and *k*_2*u*_ are their corresponding force-dependent unfolding rates which are estimated from Bell equation:

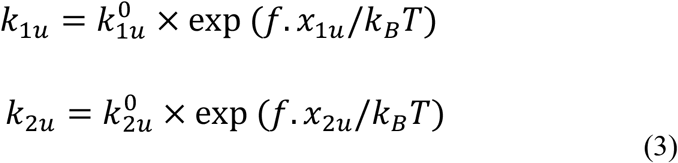

To incorporate the unfoldings from all the tip-links variants to the model, we segregated the lifetime-force data for all the step-heights ∼5, ∼12, ∼22, ∼34 nm for full-length Cdh23 EC1-27 and fitted to exponential decay (Bell model) to obtain the zero-force unfolding rate 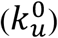 and distance to transition-state (*x*_*u*_) for the individual step height, **Figure 3g** and **Table 3**.

Further, we determined the unfolding probabilities *P*_1*u*_ and *P*_2*u*_ at each force using equations 2 and 3. Total unfolding probability is given as:

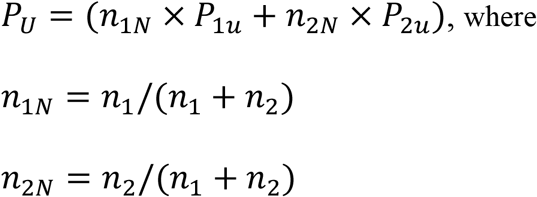

*n*_1_, *n*_2_ are the force-dependent number of unfoldings for 5 nm and 12 nm respectively, observed from the experimental data for all the tip-links variants.

Total unfolding-unbinding probability can be given as:

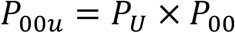

Finally, survival probability was determined as before, 1 − *P*_00*u*_

### Molecular dynamics simulations

We used the crystal structures of the complexes of EC12 domains of Cdh23 and Pcdh15 (PDB ID 4AQ8)(Sotomayor et al., 2012) and EC1-2 domains of Cdh23 and EC13 domains of Pcdh15 (PDB ID 6N2E)(Choudhary et al., 2020) as the starting structure for MD simulations. For both the complexes, the simulation setup was made with Charmm-GUI(Jo et al., 2008; Lee et al., 2016) using Charmm36 force field(Huang et al., 2017) along with TIP3P water model(Jorgensen, 1981). The long axes of the complexes were aligned to the X-axis of the box. EC12 complex was immersed in a water box with dimensions 15 × 7.2 × 7.2 nm, while EC12-EC13 complex was immersed in a box with dimensions 19 × 11 × 11 nm. For both the systems, all box angles were set equal to 90°. The system was neutralized, and the final salt concentration set to 0.15 M using sodium and chloride ions. The simulations were run using GROMACS 2020.4(Abraham et al., 2015) patched with PLUMED 2.7.4(Bonomi et al., 2009; Tribello et al., 2014). We used periodic boundary conditions and restrained the bonds involving hydrogen atoms using LINCS algorithm(Hess et al., 1997). Particle mesh Ewald method(Darden et al., 1993) was used for calculating electrostatic interactions with a cutoff of 1.2 nm for long range interactions. The system was minimized for 10,000 steps using steepest descent algorithm followed by NVT equilibration for 5 ns with restraints on the protein backbone. This was followed by FISST (Infinite Switch Simulated Tempering in Force)(Hartmann et al., 2020) simulations under NPT ensemble. We used the X-component of distance between the C-terminal residues of Cdh23 and Pcdh15 as the collective variable for EC12 complex simulations. For EC12-EC13 complex, we used the mean of the X-component of distances between the C-terminal residues of Cdh23 and Pcdh1 for the two units of the dimer. We used a force range of 10 to 150 pN with 61 interpolated points. The temperature was maintained at 303.15 K using a Nose-Hoover thermostat and for NPT simulations, the pressure was maintained at 1 bar using Parrinello-Rahman barostat.

The end-to-end distance was calculated using PLUMED and the buried surface area was calculated using sasa module of VMD. The weighted histograms for distance and buried surface area were generated using the scripts available with the PLUMED-NEST entry (plumID:20.017). The frames corresponding to different force ranges were separated using python scripts employing MDAnalysis(Michaud-Agrawal et al., 2011). The inter-protein interactions including salt-bridges, hydrogen bonds and hydrophobic interactions were calculated using PyInteraph(Tiberti et al., 2014). The pairwise residue forces were calculated using gromacs-fda(Costescu and Gräter, 2013) (release 2020) and written as signed scalars calculated by taking the norm of the force vector with the sign of the force determined based on the cosine of the angle between the force vector and vector joining the centers of masses of the two residues.

### Langevin dynamics simulations with the semiflexible filaments

We model a semiflexible filament as made up of N beads, where consecutive beads at positions *r*_*i*_ interact via the stretching energy

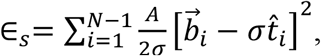

characterized by the stiffness A and bond-length *σ*. The i^th^ bond 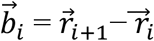 is oriented along the local tangent vector 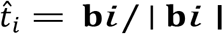. The bending rigidity κ of the semiflexible filament leads to a bending energy cost between consecutive tangent vectors(Gupta et al., 2019),

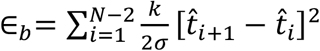

The self-avoidance of the filament is implemented through a short-ranged Weeks-Chandler-Anderson (WCA) repulsion between all the non-bonded pair of beads i and j(Shee et al., 2021),

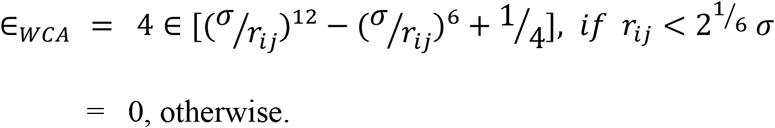

Thus, the full polymer model is described by the energy cost ∈=∈_*s*_+∈_*b*_+∈_WCA_. Bonds between two semiflexible filaments are modelled via elastic springs with the extension Δ*r* of the bond between the monomers of the two filaments generating an elastic load *f*_*i*_ = −*k*_*m*_Δ*r* with *k*_*m*_ being the stiffness of the spring.

We consider two arrangements of semiflexible filaments with elastic interactions between them as follows:

1. In one arrangement which we call a dimeric configuration, we consider two filaments with partial bonding between them as shown in Figure 4d. One end of one of the filaments is fixed. For the other filament, a constant external force ***F***_*e*_ is applied to one end of the filament. This external force can lead to the unbinding of the filaments with the bond detachment rate *ω*_*off*_. The detachment rate of individual bonds between filaments is modelled as catch-slip bond with the rate given as(Novikova and Storm, 2013)

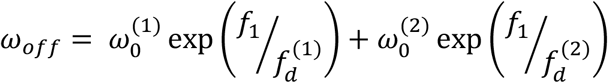 Where 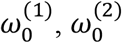 are bare detachment rates, *f* =∣ f ∣ and 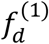 and 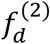 set the scales of the detachment forces. This choice of *ω*_*off*_ leads to a slip-catch-slip behavior as shown in **Figure 4d**.
2. In the second arrangement, we consider a repeated dimeric arrangement leading to a tetrameric configuration consisting of four semiflexible filaments as shown in **Figure 4e**. Filaments which are not part of the same dimeric configuration can also bind via elastic springs with spring constant *k*_*m*_. The external force now acts simultaenously on the ends of the two filaments of each of the two dimeric configurations.

We perform molecular dynamics simulations of the bead spring polymers with beads of unit mass m = 1, in the presence of a Langevin heat bath. The bath fixes the ambient temperature *k*_*B*_*T* through a Gaussian white noise which obeys ⟨*η*_*i*_ (*t*) = 0⟩ and ⟨*η*_*i*_ (*t*)*η*_*j*_(*t*^′^)⟩ = 2*αK*_*B*_*Tδ*_*ij*_ *δ*(*t* − *t*^′^), with *α* denoting the viscosity of the environment. We express energy in units of *k*_*B*_*T* = ^1^/_*β*_ lengths in units of *σ* and time in units of *τ* = *ασ*^2^/*k*_*B*_*T* = *βασ*^2^. A time step of *δt* = 0.001*τ* is used for the Langevin dynamics. We present results with averages performed over 100 initial conditions.

We use a bond stiffness of *A* = 5 × 10^3^ *k*_*B*_*T*/ *σ* for a filament of N = 35 beads which ensures smaller bond fluctuations. For an equilibrium worm like chain, the ratio of the contour length *L* = (*N* − 1)*σ* to the persistence length 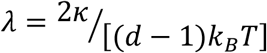 given by the rigidity parameter *u* = ^*L*^⁄_*λ*_, determines the semiflexibility of the chain with *d* as the dimension. In two dimensions, 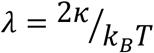. The end-to-end distribution of such a filament behaves like a completely flexible polymer with a single peak at zero separation at *u* ≈ 10 and a rigid-rod behavior with a single peak near full extension at *u* ≈ 1. We choose *u* = 3.33 where the free energy is known to show a characteristic double minimum corresponding to co-existence of both rigid rod and flexible chain behaviors. This corresponds to *κβ*/*σ* = 5.1 in the reduced units. We use *k*_*m*_ = 3 × 10^−3^ ^*A*^⁄_*σ*_ and 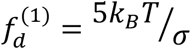 and 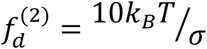. The dimensionless bare detachment rates used are 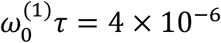 and 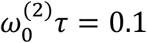.

In real units, we choose the bond length *σ* = 10*nm* = 0.01*μm*. At ambient temperature, *k*_*B*_*T* = 0.0042*pNμm*. This leads to a force unit 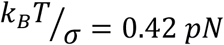. In a viscous environment with *γ* = 1*pNs*/*μm*^2^, the viscous drag on a bond can be calculated as *α* = 3*πγσ* ≈ 0.1*pNs*/*μm*. This sets the time scale 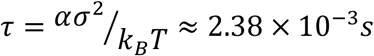. Bending rigidity *κ* = 5.1*k*_*B*_*T σ* ≈ 2.142 × 10^−4^*pNμm*^2^. *A* = 5 × 10^3^ *k*_*B*_*T*/ *σ* ≈ 2.1 × 10^3^*pN* and stiffness of the elastic bonds between filaments 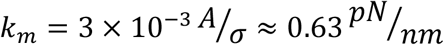. The values of the detachment forces are 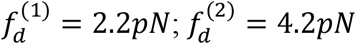 and those of the bare detachment rates are 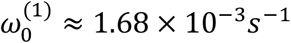 and 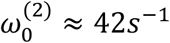 respectively.

## Acknowledgments

This work was supported by the Wellcome Trust/ DBT Indian Alliance fellowship [grant number: IA/I/15/1/501817] awarded to S. Rakshit.

We thank Professor Raj Ladher, National Centre for Biological Science, India for providing Cdh23 and Pcdh15 mammalian constructs. We thank Anuj Kumar, Dr. Amin Sagar, Sayan Das, and Gaurav Kumar Bhati for cloning of Cdh23 constructs. We thank Veerpal Kaur to perform SDS-PAGE for dimer proteins.

S. Rakshit acknowledges the financial support provided by The Wellcome Trust/DBT Intermediate fellowship by Indian Alliance and Indian Institute of Science Education and Research Mohali, India (IISERM). N.A. is thankful to CSIR-India for providing fellowship. J.P.H., S. Roy, G.K.B., are thankful to IISERM for the financial support.

## Competing interests

The authors declare no competing interests.

## Author Contributions

S. Rakshit conceived the idea. N.A., J.P.H., S.G., and S. Rakshit designed all the AFM experiments and analyzed the data. N.A expressed and purified all the monomeric proteins.

G.K.B. cloned, expressed and purified the dimer proteins. A.S. performed all the SMD simulations. S. Roy and A.C. performed all the Langevin simulations. N.A., J.P.H., A.S., and S. Roy made the figures. S. Rakshit, N.A., J.P.H., A.S., and A.C. wrote the manuscript. N.A., J.P.H. A.S., A.C., S.G., and S. Rakshit read and edited the manuscript.

## Notes

### Competing Interest Statement

The authors have declared no competing interest.

## References

Abraham, M.J., Murtola, T., Schulz, R., Páll, S., Smith, J.C., Hess, B., and Lindah, E. (2015). Gromacs: High performance molecular simulations through multi-level parallelism from laptops to supercomputers. SoftwareX 1–2, 19–25. https://doi.org/10.1016/j.softx.2015.06.001.

Ahmed, Z.M., Riazuddin, S., Aye, S., Ali, R.A., Venselaar, H., Belyantseva, P.P., Qasim, M., Riazuddin, S., and B, T. (2008). Gene structure and mutant alleles of PCDH15: nonsyndromic deafness DFNB23 and type 1 Usher syndrome. Hum. Genet. 124, 215–223. https://doi.org/10.1007/s00439-008-0543-3.

Araya-Secchi, R., Neel, B.L., and Sotomayor, M. (2016). An elastic element in the protocadherin-15 tip link of the inner ear. Nat. Commun. 7, 1–14. https://doi.org/10.1038/ncomms13458.

Arora, N., Hazra, J.P., and Rakshit, S. (2021). Anisotropy in mechanical unfolding of protein upon partner-assisted pulling and handle-assisted pulling. Commun. Biol. 4, 1–10. https://doi.org/10.1038/s42003-021-02445-y.

Astuto, L.M., Bork, J.M., Weston, M.D., Askew, J.W., Fields, R.R., Orten, D.J., Ohliger, S.J., Riazuddin, S., Morell, R.J., Khan, S., et al. (2002). CDH23 Mutation and Phenotype Heterogeneity: A Profile of 107 Diverse Families with Usher Syndrome and Nonsyndromic Deafness. Am. J. Hum. Genet. 71, 262–275. https://doi.org/10.1086/341558.

Bartsch, T.F., Hengel, F.E., Oswald, A., Dionne, G., and Chipendo, I. V (2018). The elasticity of individual protocadherin 15 molecules implicates cadherins as the gating springs for hearing. Proc. Natl. Acad. Sci. 116, 11048–11056. https://doi.org/10.1101/503029.

Basu, A., Lagier, S., Vologodskaia, M., Fabella, B.A., and Hudspeth, A.J. (2016). Direct mechanical stimulation of tip links in hair cells through DNA tethers. Elife 5:e16041. 1–10. https://doi.org/10.7554/eLife.16041.

Bell, G.I. (1978). Models for the Specific Adhesion of Cells to Cells. Science (80-.). 200, 618–627. https://doi.org/10.1017/s0001867800035254.

Bohbot-Raviv, Y., Zhao, W.Z., Feingold, M., Wiggins, C.H., and Granek, R. (2004). Relaxation dynamics of semiflexible polymers. Phys. Rev. Lett. 92, 1–4. https://doi.org/10.1103/PhysRevLett.92.098101.

Bonomi, M., Branduardi, D., Bussi, G., Camilloni, C., Provasi, D., Raiteri, P., Donadio, D., Marinelli, F., Pietrucci, F., Broglia, R.A., et al. (2009). PLUMED: A portable plugin for free-energy calculations with molecular dynamics. Comput. Phys. Commun. 180, 1961–1972. https://doi.org/10.1016/j.cpc.2009.05.011.

Brujic, J., Hermans, R.I.Z., Garcia-Manyes, S., Walther, K.A., and Fernandez, J.M. (2007). Dwell-time distribution analysis of polyprotein unfolding using force-clamp spectroscopy. Biophys. J. 92, 2896–2903. https://doi.org/10.1529/biophysj.106.099481.

Cao, Y., Kuske, R., and Li, H. (2008). Direct observation of Markovian behavior of the mechanical unfolding of individual proteins. Biophys. J. 95, 782–788. https://doi.org/10.1529/biophysj.107.128298.

Choudhary, D., Narui, Y., Neel, B.L., Wimalasena, L.N., Klanseck, C.F., De-La-Torre, P., Chen, C., Araya-Secchi, R., Tamilselvan, E., and Sotomayor, M. (2020). Structural determinants of protocadherin-15 mechanics and function in hearing and balance perception. Proc. Natl. Acad. Sci. U. S. A. 117, 24837–24848. https://doi.org/10.1073/pnas.1920444117.

Corey, D.P., and Hudspeth, A.J. (1983). Kinetics of the receptor current in bullfrog saccular hair cells. J. Neurosci. 3, 962–976..

Costescu, B.I., and Gräter, F. (2013). Time-resolved force distribution analysis. BMC Biophys. 6, 1–5. https://doi.org/10.1186/2046-1682-6-5.

Darden, T., York, D., and Pedersen, L. (1993). Particle mesh Ewald: An N·log(N) method for Ewald sums in large systems. J. Chem. Phys. 98, 1–5. https://doi.org/10.1063/1.464397.

Dembo, M., Torney, D.C., Saxman, K., and Hammer, D. (1988). The reaction-limited kinetics of membrane-to-surface adhesion and detachment. Proc. R. Soc. B Biol. Sci. 55–83. https://doi.org/10.1098/rspb.1988.0038.

Gibson, D.G., Young, L., Chuang, R.Y., Venter, J.C., Hutchison, C.A., and Smith, H.O. (2009). Enzymatic assembly of DNA molecules up to several hundred kilobases. Nat. Methods 6, 343–345. https://doi.org/10.1038/nmeth.1318.

Granzier, H., Kellermayer, M., Helmes, M., and Trombitás, K. (1997). Titin elasticity and mechanism of passive force development in rat cardiac myocytes probed by thin-filament extraction. Biophys. J. 73, 2043–2053. https://doi.org/10.1016/S0006-3495(97)78234-1.

Gupta, N., Chaudhuri, A., and Chaudhuri, D. (2019). Morphological and dynamical properties of semiflexible filaments driven by molecular motors. Phys. Rev. E 99, 1–10. https://doi.org/10.1103/PhysRevE.99.042405.

Hartmann, M.J., Singh, Y., Vanden-Eijnden, E., and Hocky, G.M. (2020). Infinite switch simulated tempering in force (FISST). J. Chem. Phys. 152, 1–10. https://doi.org/10.1063/5.0009280.

Hazra, J.P., Sagar, A., Arora, N., Deb, D., Kaur, S., and Rakshit, S. (2019). Broken force dispersal network in tip-links by the mutations induces hearing-loss. Biochem. J. 476, 2411–2425. https://doi.org/10.1101/614610.

Hess, B., Bekker, H., Berendsen, H.J.C., and Fraaije, J.G.E.M. (1997). LINCS: A Linear Constraint Solver for molecular simulations. J. Comput. Chem. 18, 1463–1472. https://doi.org/10.1002/(SICI)1096-987X(199709)18:12<1463::AID-JCC4>3.0.CO;2-H.

Honig, B., Liang, X., Shapiro, L., Potter, C.S., Dionne, G., Rapp, M., Qiu, X., Carragher, B., Ahlsen, G., Peng, G., et al. (2018). Mechanotransduction by PCDH15 Relies on a Novel cis-Dimeric Architecture. Neuron 99, 480–492. https://doi.org/10.1016/j.neuron.2018.07.006.

Huang, J., Rauscher, S., Nawrocki, G., Ran, T., Feig, M., De Groot, B.L., Grubmüller, H., and MacKerell, A.D. (2017). CHARMM36m: An improved force field for folded and intrinsically disordered proteins. Nat. Methods 14, 71–73. https://doi.org/10.1038/nmeth.4067.

Hudspeth, A.J. (1989). How the ear’s works work. Nature 341, 397–404. https://doi.org/10.1038/341397a0.

Hudspeth, A.J. (1992). Hair-bundle mechanics and a model for mechanoelectrical transduction by hair cells. Soc. Gen. Physiol. Ser. 47, 357–370..

Hudspeth, A.J. (1997). How hearing happens. Neuron 947–950. https://doi.org/10.1016/S0896-6273(00)80385-2.

Hutter, J.L., and Bechhoefer, J. (1993). Calibration of atomic-force microscope tips. Rev. Sci. Instrum. 64, 1868–1873. https://doi.org/10.1063/1.1143970.

Jaiganesh, A., De-la-torre, P., Patel, A.A., Termine, D.J., Velez-cortes, F., Chen, C., and Sotomayor, M. (2018). Zooming in on Cadherin-23 : Structural Diversity and Potential Mechanisms of Inherited Deafness. Structure 26, 1–16. https://doi.org/10.1016/j.str.2018.06.003.

Jo, S., Kim, T., Iyer, V.G., and Im, W. (2008). CHARMM-GUI: A web-based graphical user interface for CHARMM. J. Comput. Chem. 29, 1–7. https://doi.org/10.1002/jcc.20945.

Jorgensen, W.L. (1981). Quantum and statistical mechanical studies of liquids. 11. Transferable intermolecular potential functions. Application to liquid methanol including internal rotation. J. Am. Chem. Soc. 103, 335–340. https://doi.org/10.1021/ja00392a017.

Kachar, B., Parakkal, M., Kurc, M., Zhao, Y., and Gillespie, P.G. (2000). High-resolution structure of hair-cell tip links. Proc. Natl. Acad. Sci. 97, 13336–13341. https://doi.org/10.1073/pnas.1311228110.

Kazmierczak, P., Milligan, R.A., Wilson-Kubalek, E.M., Tokita, J., Kachar, B., Sakaguchi, H., and Müller, U. (2007). Cadherin 23 and protocadherin 15 interact to form tip-link filaments in sensory hair cells. Nature 449, 87–91. https://doi.org/10.1038/nature06091.

Lee, G., Abdi, K., Jiang, Y., Michaely, P., Bennett, V., and Marszalek, P.E. (2006). Nanospring behaviour of ankyrin repeats. Nature 440, 246–249. https://doi.org/10.1038/nature04437.

Lee, J., Cheng, X., Swails, J.M., Yeom, M.S., Eastman, P.K., Lemkul, J.A., Wei, S., Buckner, J., Jeong, J.C., Qi, Y., et al. (2016). CHARMM-GUI Input Generator for NAMD, GROMACS, AMBER, OpenMM, and CHARMM/OpenMM Simulations Using the CHARMM36 Additive Force Field. J. Chem. Theory Comput. 12, 405–413. https://doi.org/10.1021/acs.jctc.5b00935.

Lou, J., and Zhu, C. (2007). A structure-based sliding-rebinding mechanism for catch bonds. Biophys. J. 92, 1471–1485. https://doi.org/10.1529/biophysj.106.097048.

Michaud-Agrawal, N., Denning, E.J., Woolf, T.B., and Beckstein, O. (2011). MDAnalysis: A toolkit for the analysis of molecular dynamics simulations. J. Comput. Chem. 32, 1–9. https://doi.org/10.1002/jcc.21787.

Miyagawa, M., Nishio, S. ya, and Usami, S. ichi (2012). Prevalence and clinical features of hearing loss patients with cdh23 mutations: A large cohort study. PLoS One 7, 1–12. https://doi.org/10.1371/journal.pone.0040366.

Mulhall, E.M., Ward, A., Yang, D., Koussa, M.A., Corey, D.P., and Wong, W.P. (2021). Single-molecule force spectroscopy reveals the dynamic strength of the hair-cell tip-link connection. Nat. Commun. 12, 1–15. https://doi.org/10.1038/s41467-021-21033-6.

Muller, U., and Gillespie, P.G. (2010). Mechanotransduction by Hair Cells: Models, Molecules, and Mechanisms. Cell 139, 33–44. https://doi.org/10.1016/j.cell.2009.09.010.Mechanotransduction.

Nair, A., Chandel, S., Mitra, M.K., Muhuri, S., and Chaudhuri, A. (2016). Effect of catch bonding on transport of cellular cargo by dynein motors. Phys. Rev. E 94, 1–5. https://doi.org/10.1103/PhysRevE.94.032403.

Novikova, E.A., and Storm, C. (2013). Contractile fibers and catch-bond clusters: A biological force sensor? Biophys. J. 105, 1336–1345. https://doi.org/10.1016/j.bpj.2013.07.039.

Oroz, J., Galera-Prat, A., Hervás, R., Valbuena, A., Fernández-Bravo, D., and Carrión-Vázquez, M. (2019). Nanomechanics of tip-link cadherins. Sci. Rep. 9, 1–9. https://doi.org/10.1038/s41598-019-49518-x.

Ott, W., Schoeler, C., Nash, M.A., Bernardi, R.C., Gaub, H.E., Schulten, K., Malinowska, K.H., Bayer, E.A., and Durner, E. (2015). Mapping Mechanical Force Propagation through Biomolecular Complexes. Nano Lett. 15, 7370–7376. https://doi.org/10.1021/acs.nanolett.5b02727.

Pereverzev, Y. V., Prezhdo, O. V., Forero, M., Sokurenko, E. V., and Thomas, W.E. (2005). The two-pathway model for the catch-slip transition in biological adhesion. Biophys. J. 89, 1446–1454. https://doi.org/10.1529/biophysj.105.062158.

Pickles, J.O., Comis, S.D., and Osborne, M.P. (1984a). Cross-links between stereocilia in the guinea pig organ of Corti, and their possible relation to sensory transduction. Hear. Res. 15, 103–112. https://doi.org/10.1016/0378-5955(84)90041-8.

Pickles, J.O., Comis, S.D., and Osborne, M.P. (1984b). Cross-links between stereocilia in the guinea pig organ of Corti, and their possible relation to sensory transduction. Hear. Res. 15, 103–112. https://doi.org/10.1016/0378-5955(84)90041-8.

Powers, R.E., Gaudet, R., and Sotomayor, M. (2017). A Partial Calcium-Free Linker Confers Flexibility to Inner-Ear Protocadherin-15. Structure 25, 482–495. https://doi.org/10.1016/j.str.2017.01.014.

Rakshit, S., and Sivasankar, S. (2014). Biomechanics of cell adhesion: How force regulates the lifetime of adhesive bonds at the single molecule level. Phys. Chem. Chem. Phys. 16, 2211–2223. https://doi.org/10.1039/c3cp53963f.

Rakshit, S., Zhang, Y., Manibog, K., Shafraz, O., and Sivasankar, S. (2012). Ideal, catch, and slip bonds in cadherin adhesion. Proc. Natl. Acad. Sci. U. S. A. 109, 18815–18820. https://doi.org/10.1073/pnas.1208349109.

Sahly, I., Dufour, E., Schietroma, C., Michel, V., Bahloul, A., Perfettini, I., Pepermans, E., Estivalet, A., Carette, D., Aghaie, A., et al. (2012). Localization of usher 1 proteins to the photoreceptor calyceal processes, which are absent from mice. J. Cell Biol. 199, 381–399. https://doi.org/10.1083/jcb.201202012.

Shee, A., Gupta, N., Chaudhuri, A., and Chaudhuri, D. (2021). A semiflexible polymer in a gliding assay: reentrant transition, role of turnover and activity. Soft Matter 17, 2120–2131. https://doi.org/10.1039/d0sm01181a.

Sotomayor, M., Corey, D.P., and Schulten, K. (2005). In search of the hair-cell gating spring: Elastic properties of ankyrin and cadherin repeats. Structure 13, 669–682. https://doi.org/10.1016/j.str.2005.03.001.

Sotomayor, M., Weihofen, W.A., Gaudet, R., and Corey, D.P. (2010). Structural Determinants of Cadherin-23 Function in Hearing and Deafness. Neuron 66, 85–100. https://doi.org/10.1016/j.neuron.2010.03.028.Structural.

Sotomayor, M., Weihofen, W., Gaudet, R., and Corey, D.P. (2012). Structure of a Force-Conveying Cadherin Bond Essential for Inner-Ear Mechanotransduction. Nature 492, 128–132. https://doi.org/10.1038/nature11590.

Srinivasan, S., Hazra, J.P., Singaraju, G.S., Deb, D., and Rakshit, S. (2017). ESCORTing proteins directly from whole cell-lysate for single-molecule studies. Anal. Biochem. 535, 35–42. https://doi.org/10.1016/j.ab.2017.07.022.

Thomas, W., Forero, M., Yakovenko, O., Nilsson, L., Vicini, P., Sokurenko, E., and Vogel, V. (2006). Catch-bond model derived from allostery explains force-activated bacterial adhesion. Biophys. J. 90, 753–764. https://doi.org/10.1529/biophysj.105.066548.

Tiberti, M., Invernizzi, G., Lambrughi, M., Inbar, Y., Schreiber, G., and Papaleo, E. (2014). PyInteraph: A framework for the analysis of interaction networks in structural ensembles of proteins. J. Chem. Inf. Model. 54, 1537–1551. https://doi.org/10.1021/ci400639r.

Tribello, G.A., Bonomi, M., Branduardi, D., Camilloni, C., and Bussi, G. (2014). PLUMED 2: New feathers for an old bird. Comput. Phys. Commun. 185, 604–613. https://doi.org/10.1016/j.cpc.2013.09.018.

