## Supplementary File for "Tip-links serve as force-pass filter to fulfil the role of gating-springs"

Figure S1: Hydrogen bond survival with force obtained from FISST simulations 3
Figure S2: Survival of hydrophobic interactions with force obtained from FISST simulations 4
Figure S3: Correlation coefficient between the inter-residue forces for all the intra-protein residue pairs 5
Figure S4: Unfolding prior to unbinding prolongs the bond lifetime 6
Figure S5: Percentage of unfolding events in different tip-links variants 7
Figure S6: SDS-PAGE confirmed the dimerization of the cadherin proteins in the solution 8
Figure S7: Unfolding step height distribution for heterotetrameric tip-links 9
Figure S8: Resulting amplitudes A1, and A2 from the double-exponential fitting of the survival plots 10
Figure S9: Elastic nature of 5nm extension from the SMD simulations and proposed spring-dashpot model for of gating-spring 11
Figure S10: Force-dependent lifetime behaviour of semiflexible filaments linked with only slip-bonds 12

Supplementary Movie 1 13
Supplementary Movie 2 13

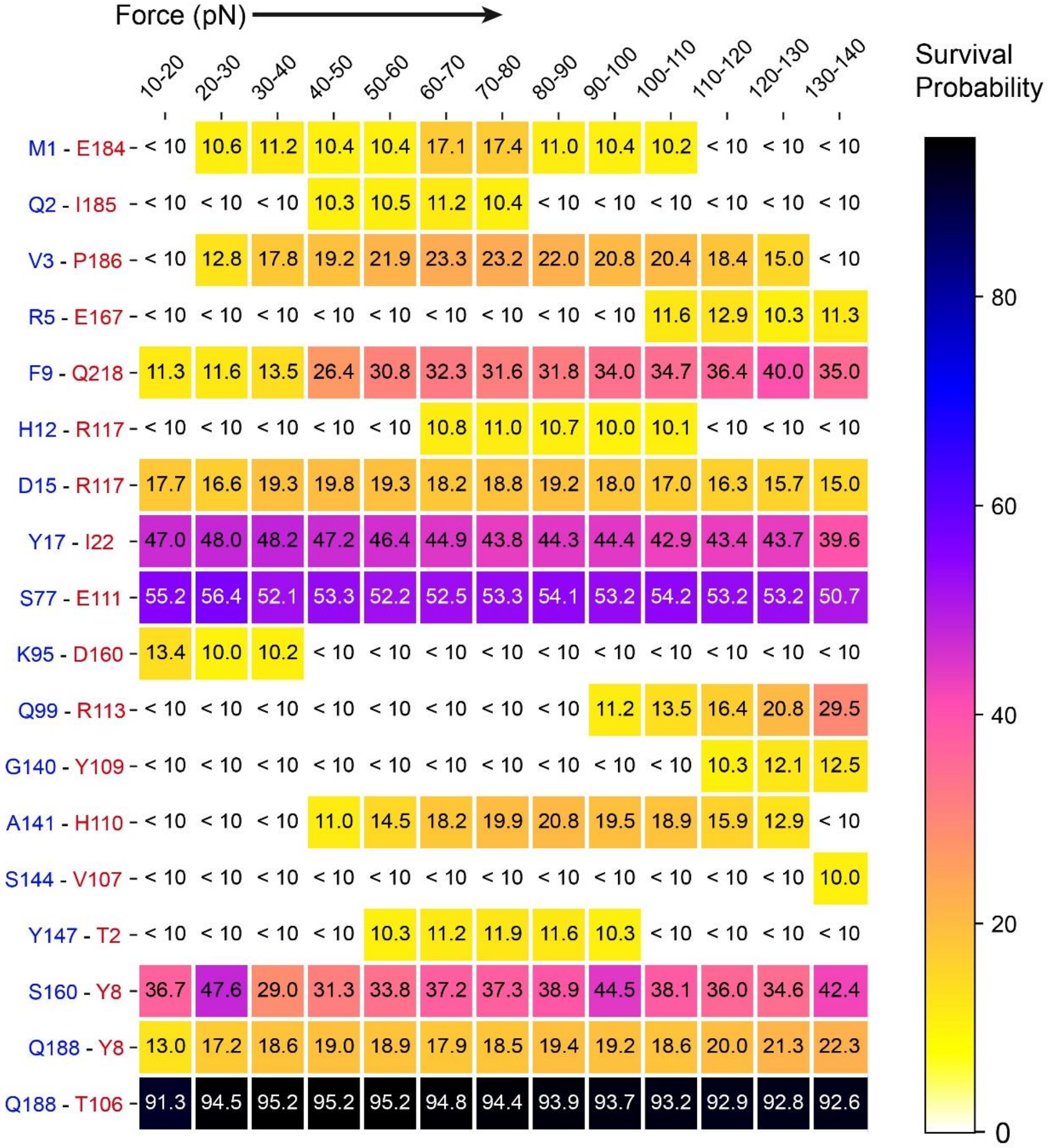


**Supplementary Figure S1: Hydrogen bond survival with force obtained from FISST simulations (in support of figure 2).** The percentage of frames showing the existence of specified hydrogen-bond interactions at different force ranges. The residue pair on the left depicts the hydrogen bond between residues of Cdh23 (blue) and Pcdh15 (red).

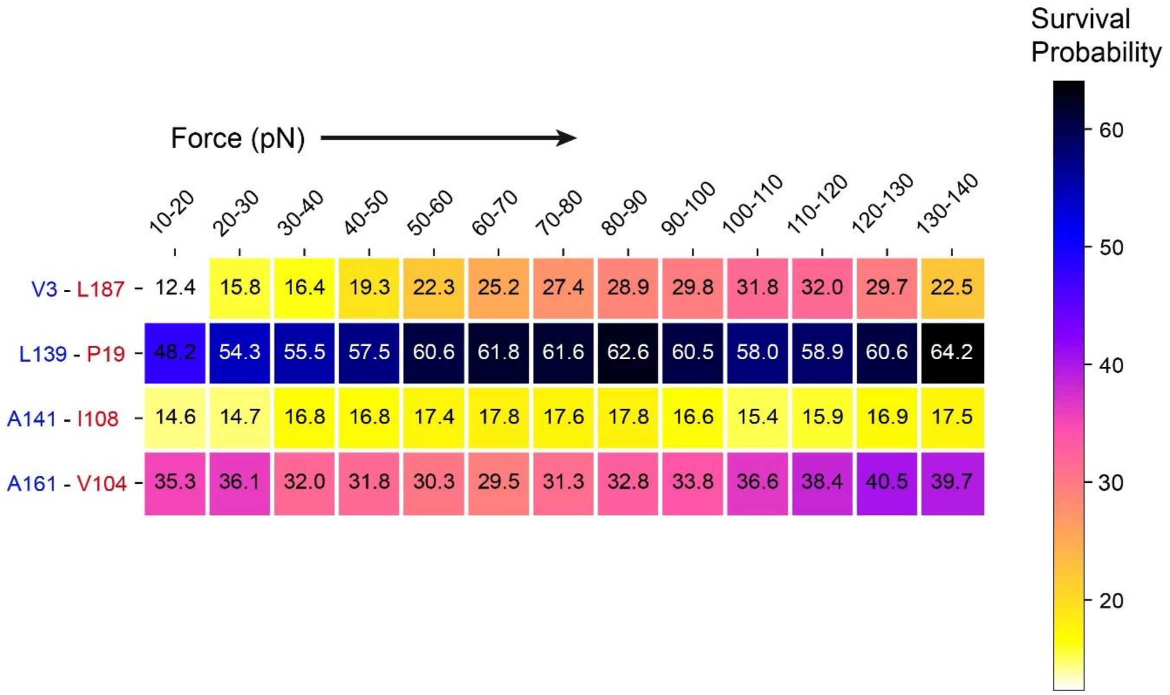


**Supplementary Figure S2: Survival of hydrophobic interactions with force obtained from FISST simulations (in support of figure 2).** The percentage of frames showing the existence of specified hydrophobic interactions at different force ranges. The residue pair on the left depicts the hydrophobic interaction between residues of Cdh23 (blue) and Pcdh15 (red).

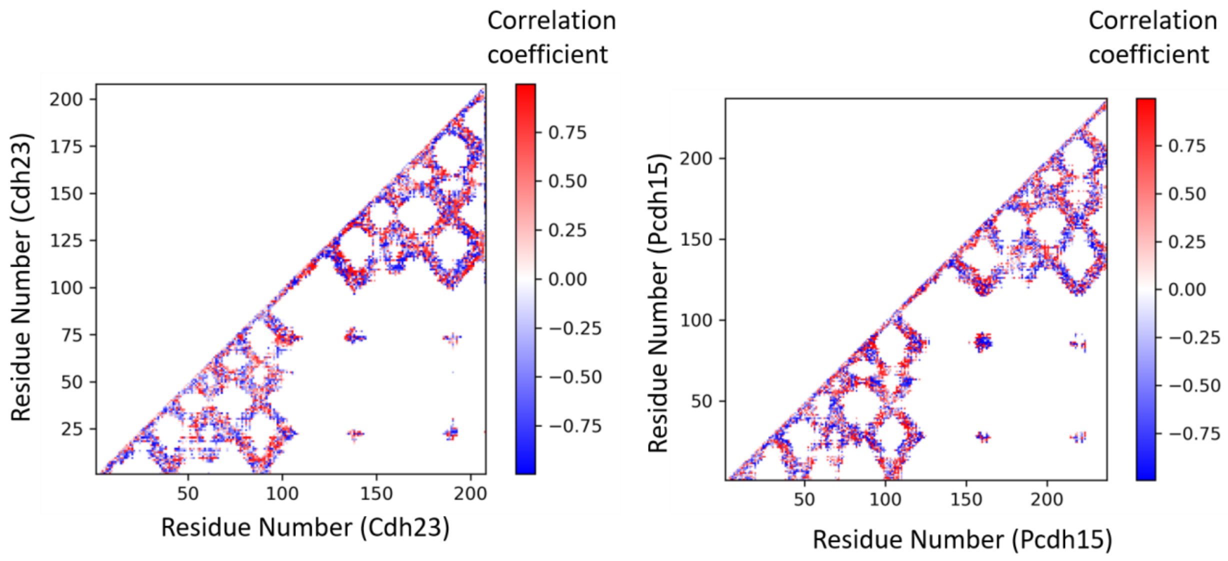


**Supplementary Figure S3: Correlation coefficient between the inter-residue forces and the applied force ($r^f_{af}$) for all the intra-protein residue pairs of Cdh23 and Pcdh15 EC1-2 (in support of figure 2).**

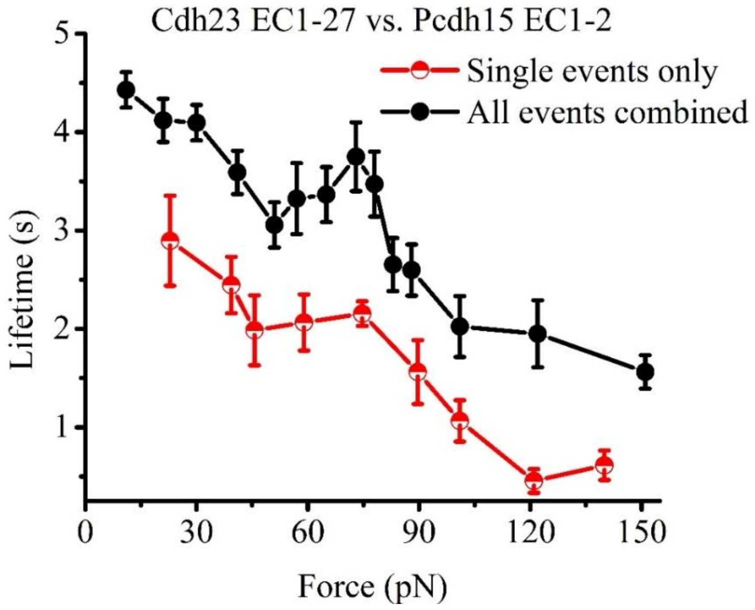


**Supplementary Figure S4. Unfolding prior to unbinding prolongs the bond lifetime (in support of figure 3).** Unbinding events without undergoing any unfolding registered a lower overall lifetime (red) compared to the events which showed unfolding prior to unbinding (black) for Cdh23 EC1-27 vs Pcdh15 EC1-2 interaction.

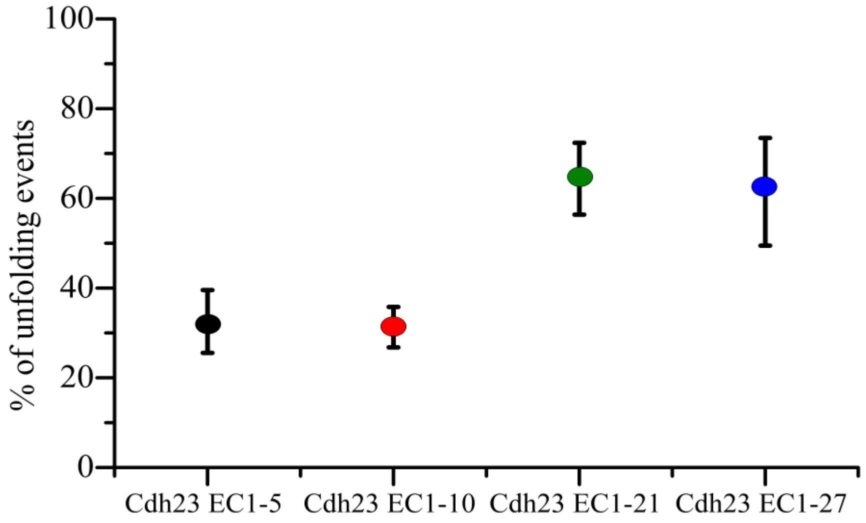


**Supplementary Figure S5. Percentage of unfolding events obtained in the force-clamp measurements for different tip-links variants. (In support of figure 3).** % of force-clamp events undergoing unfolding before unbinding increases with increasing the EC domain numbers.

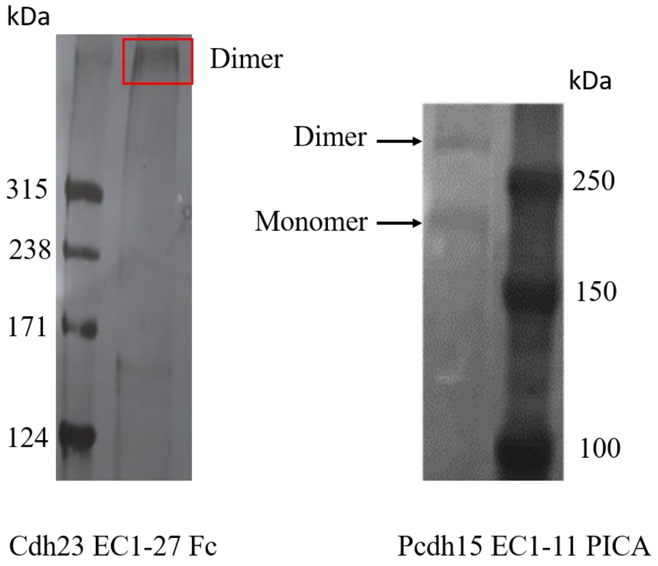


**Supplementary Figure S6. SDS-PAGE followed by silver-staining confirmed the dimerization of the individual tip-links cadherin proteins in the solution (in support of figure 4).** The left panel shows the silver-stained gel for Cdh23 EC1-27 Fc protein at a higher molecular weight corresponding to the dimer. The right panel shows the monomer and dimer bands of Pcdh15 EC1-11 PICA in a silver-stained SDS-PAGE gel.

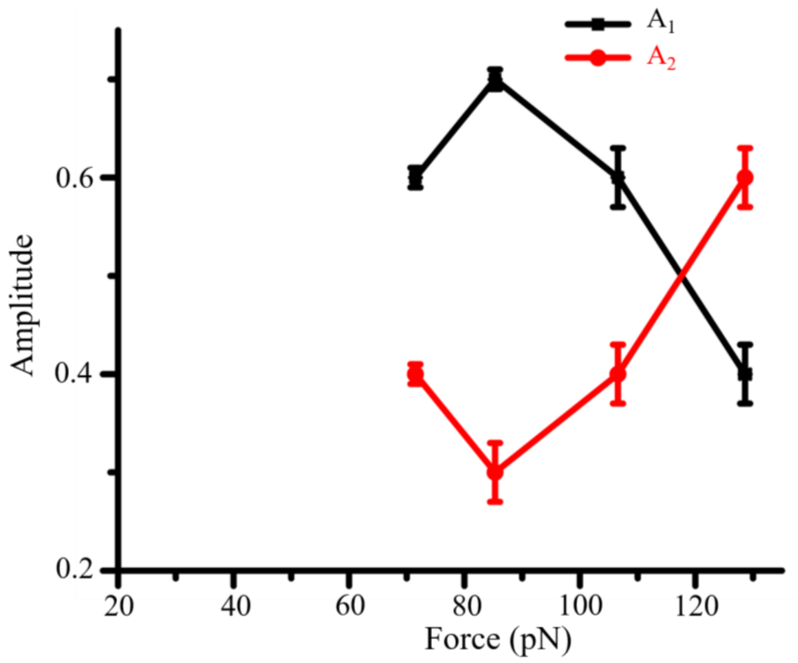


**Supplementary Figure S7. Resulting amplitudes $A_1$, and $A_2$ from the double-exponential fitting of the survival plots for heterotetrameric tip-links complex at higher forces (>70 pN) (in support of figure 4).** Amplitude corresponding to the higher-lifetime component (black) decreases with force whereas corresponding to the lower lifetime component (red) increases with force. Errors are the standard errors obtained from the exponential fitting of the survival probability curves.

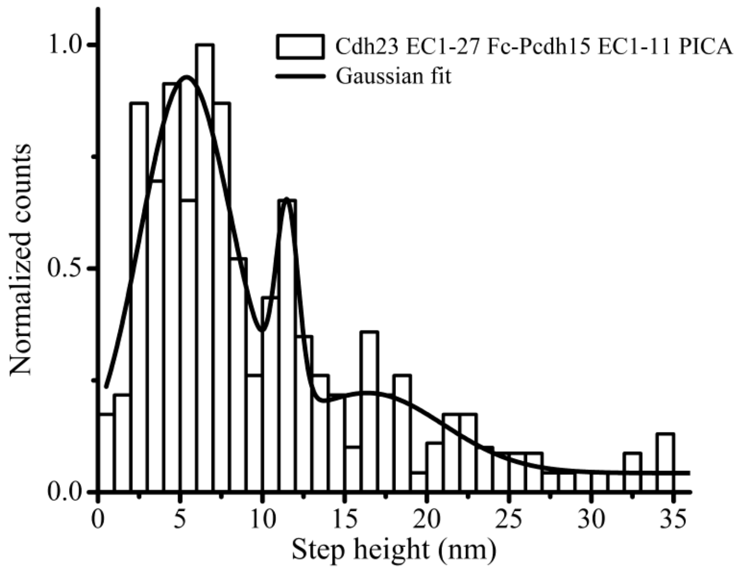


**Supplementary Figure S8. Unfolding step height distribution for heterotetrameric tip-links Cdh23 EC1-27 Fc-Pcdh15 EC1-11 PICA (in support of figure 4).** Gaussian fitting of step height distribution resulted in three major peaks at 5.3±0.3, 11.5±0.2, and 16.4±2.8 nm.

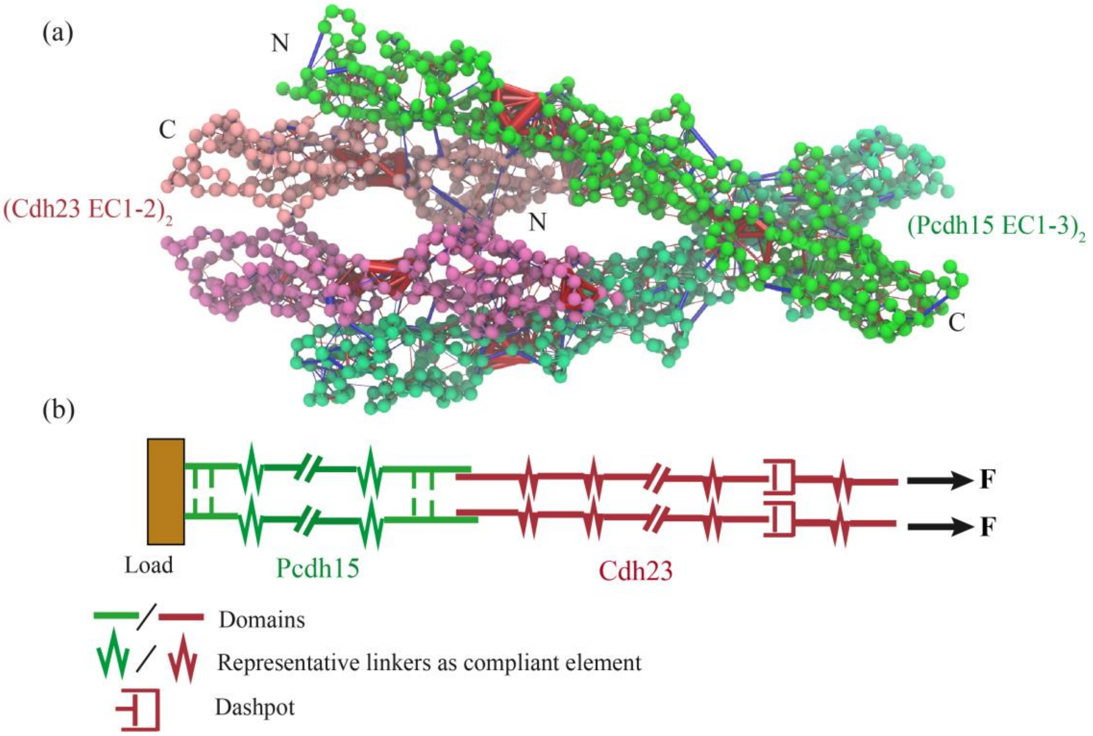


**Supplementary Figure S9. Elastic nature of 5nm extension from the SMD simulations and proposed spring-dashpot model for tip-links as gating-spring**. **(in support of figure 4). (a)** Force-distribution analysis for the dimer of Cdh23 EC1-2-Pcdh15 EC1-3 indicates that the linker between the EC1-2 and EC2-3 are pre-compressed (red cylinders) and thus, imparts mechanical stability to the protein against force. **(b)** Schematic depiction of tip-links in the spring-dashpot model. Here, Pcdh15(green), represented with a series of springs, is connected to load at C-terminus, and Cdh23 (red) is represented with springs and dashpots in series. Dashpots are relatively less in number representing the elongation of ~12 nm or higher presumably non-elastically by the auditory inputs.

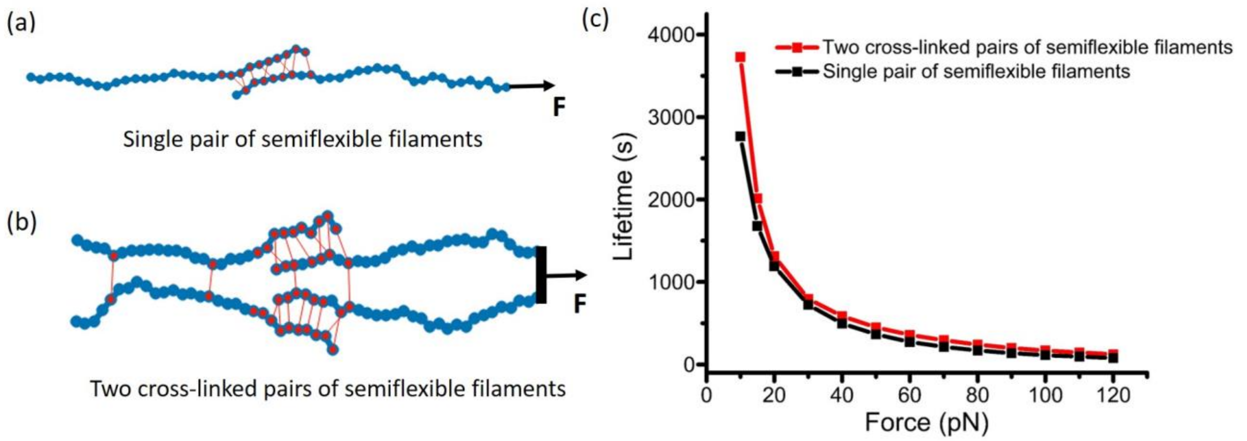


**Supplementary Figure S10: Force-dependent lifetime behaviour of semiflexible filaments linked with only slip-bonds (in support of figure 4). (a and b)** Schematic representation of single and double pair of semiflexible filaments coupled with multiple elastic slip-bonds. **(c)** This slip-bonded arrangement between chains resulted in exclusive slip-bond with force for both systems.

**Supplementary Movie 1**

The dimeric arrangement of two semiflexible filaments, partially attached to one another via elastic bonds (red) with catch-slip dissociation characteristics. The bonds dissociate in the presence of elastic load when one of the filaments is pulled externally.

**Supplementary Movie 2**

The tetrameric arrangement, where the two filaments from each of the dimeric arrangements are pulled simultaneously with the external force at one end. Additionally, the anchored filaments are cross-linked resembling the cis-dimers of Pcdh15. Also, we cross-linked the polymer couples that are pulled, at their cross-linking interface. This polymer couple resembles Cdh23. We observed the rebinding of the bonds in this tetrameric arrangement before the complete unbinding.
